# The ABCE1 capsid assembly pathway is conserved between primate lentiviruses and the non-primate lentivirus feline immunodeficiency virus

**DOI:** 10.1101/183848

**Authors:** Jonathan C. Reed, Nick Westergreen, Brook C. Barajas, Dylan Ressler, Daryl Phuong, John V. Swain, Vishwanath R. Lingappa, Jaisri R. Lingappa

## Abstract

During immature capsid assembly in cells, the Gag protein of HIV-1 and other primate lentiviruses co-opts a host RNA granule, forming a pathway of assembly intermediates that contains host components, including two cellular enzymes shown to facilitate assembly, ABCE1 and DDX6. Here we asked whether a non-primate lentivirus, feline immunodeficiency virus (FIV), also forms such RNA-granule-derived intracellular capsid assembly intermediates. First, we found that, unlike for HIV-1, the FIV completed immature capsid and the largest putative assembly intermediate are unstable during analysis. Next, we identified *in situ* cross-linking conditions that overcame this problem and revealed the presence of FIV Gag complexes that correspond in size to early and late HIV-1 assembly intermediates. Because assembly-defective HIV-1 Gag mutants are arrested at specific intracellular assembly intermediates, we asked if a similar arrest is also observed for FIV. We analyzed four FIV Gag mutants, including three not previously studied that we identified based on sequence and structural similarity to HIV-1 Gag, and found that each is assembly-defective and arrested at the same intermediate as the corresponding HIV-1 mutant. Further evidence that these FIV Gag-containing complexes correspond to assembly intermediates came from co-immunoprecipitation studies demonstrating that FIV Gag is associated with ABCE1 and DDX6, as shown previously for HIV-1. Finally, we validated these co-immunoprecipitations with a proximity ligation assay that revealed co-localization between assembly-competent FIV Gag and ABCE1 *in situ*. Together, these data offer novel structure-function insights and indicate that primate and non-primate lentiviruses form intracellular capsid assembly intermediates derived from ABCE1-containing RNA granules.

**Importance:** Like HIV-1, FIV Gag assembles into immature capsids; however, it is not known whether FIV Gag progresses through a pathway of immature capsid assembly intermediates derived from host RNA granules, as shown for HIV-1 Gag. Here we asked whether FIV Gag forms complexes similar in size to HIV-1 assembly intermediates and if FIV Gag is associated with ABCE1 and DDX6, two host enzymes that facilitate HIV-1 immature capsid assembly that are found in HIV-1 assembly intermediates. Our studies identified FIV Gag-containing complexes that closely resemble HIV-1 capsid assembly intermediates, showed that known and novel assembly-defective FIV Gag mutants fail to progress past these putative intermediates, and utilized biochemical and imaging approaches to demonstrate association of FIV Gag with ABCE1 and DDX6. Thus, we conclude that viral-host interactions important for immature capsid assembly are conserved between primate and non-primate lentiviruses, and could yield important targets for future antiviral strategies.

## Introduction

The feline immunodeficiency virus (FIV) is one of the most common infectious diseases in domestic cats with a prevalence in North America of ~2%. The natural history of FIV infection follows a pattern similar to the human immunodeficiency virus type 1 (HIV-1) starting with an initial acute infection phase, followed by an asymptomatic phase of variable length, and a terminal phase that results in feline acquired immunodeficiency syndrome (feline AIDS)(reviewed in (1)). A commercial FIV vaccine has been developed for cats, but it may not provide complete protection from circulating FIV strains (2, 3). In infected cats, when less aggressive management fails to prevent recurrent disease, antiretroviral chemotherapy can be initiated using antiretroviral drugs. The antiretroviral drug typically prescribed is the HIV-1 reverse transcriptase inhibitor Zidovudine (AZT) (4). Although AZT can improve quality of life and extend life expectancy, resistance can develop quickly and treatment can have serious side effects (reviewed in (5)). While most infected cats do not have severe symptoms, given that cats can live healthier lives with treatment, drugs that specifically and potently inhibit FIV replication could be of benefit. In addition, since FIV infection of cats can result in AIDS, it has been proposed that such infection could serve as a useful model system for studying new antiviral treatments and vaccines for HIV-1.

To utilize FIV infection as a model for HIV-1 infection would require knowing the similarities and differences between the two, which have been defined for some stages of the two viral life cycles (reviewed in (6)). FIV has some notable differences relative to HIV-1; for example FIV encodes a dUTPase, which is not found in the primate lentiviruses, and FIV encodes fewer accessory proteins compared to HIV-1. Additionally FIV encodes OrfA, which may not have a direct HIV-1 ortholog. Here we were interested in addressing whether mechanisms involved in intracellular immature capsid assembly are conserved between HIV-1 and FIV. Like all exogenous retroviruses, HIV-1 and FIV, encode three proteins that carry out essential steps common to all retroviral lifecycles - Gag, Pol, and Env. Gag is the only viral protein required for immature capsid assembly. The domains of the Gag polyprotein required for immature capsid assembly have been studied extensively for HIV-1 (reviewed in (7)). The HIV-1 matrix domain (MA) confers plasma membrane targeting, and the HIV-1 capsid domain (CA), which contains an N-terminal and C-terminal subdomain (CA-NTD and CA-CTD, respectively), provides important Gag-Gag contacts required for the immature capsid structure. The HIV-1 nucleocapsid domain (NC) mediates both specific interactions with viral genomic RNA and non-specific RNA interactions, while the HIV-1 late domain (p6) recruits host factors important for budding the virus from infected cells. FIV Gag encodes the same domains as HIV-1 Gag and mutational studies have confirmed that the FIV Gag domains (MA, CA, NC, and late domain p2) function similarly to their HIV-1 counterparts (8–11). For example, the late domain of both viruses encodes a TSG101 binding motif to recruit TSG101 and ESCRT machinery to promote budding (11). In both cases, full-length Gag in the immature virus undergoes maturation when the viral protease cleaves Gag into separate domains (MA, CA, p1, NC, and p2 for FIV; MA, CA, sp1, NC, sp2, and p6 for HIV-1)(12). Importantly, numerous assembly-defective mutants have been generated in structure-function analyses of HIV-1 Gag (reviewed in (13)), but are lacking for FIV Gag. Thus, one goal of our study was to generate a diverse set of assembly-defective FIV Gag point mutants that would serve as tools for studying the mechanism of FIV Gag assembly.

If the HIV-1 and FIV Gag proteins are similar, the likelihood of shared intracellular mechanisms of assembly would be high. At the level of overall amino acid sequence, conservation among different retroviral Gag proteins is low, with the major homology region of Gag being a notable exception (14–16). However, known atomic structures of CA subdomains from different retroviruses display a high degree of conservation (17–19). Thus, conservation of overall CA structure between different retroviruses argues for shared mechanisms for immature capsid assembly. This could include conservation of specific Gag-Gag interactions in immature capsids, but also conservation of Gag-host interactions that may facilitate immature capsid assembly by common mechanisms within cells. To determine whether FIV and HIV-1 Gag assemble via similar mechanisms, here we asked whether intracellular events that occur during HIV-1 immature capsid assembly are conserved for FIV.

HIV-1 immature capsid assembly proceeds via a sequential pathway of post-translational assembly intermediates (20–22). At steady state, four major intermediates of increasing size can be identified in cells, starting with the first intermediate (~10S), and followed by progression through the subsequent ~80S, ~150S, and ~500S intermediates to the final completed immature capsid (~750S) (20–22). Evidence that the complexes observed at steady-state are *bona fide* assembly intermediates come from pulse-chase studies showing that they are formed sequentially (20, 21), and mutational analyses showing that all assembly-defective HIV-1 Gag mutants studied to date are arrested at different steps in this assembly pathway (22–25). Moreover, other components found in the released virus, such as the HIV-1 Vif protein, GagPol, and genomic RNA are also found in assembly intermediates (21, 22, 26). Notably, host proteins have been shown to play a role in facilitating these events. Immunodepletion studies from cell-free extracts and dominant negative studies in cells established that the cellular enzyme ATP binding cassette protein E1 (ABCE1) facilitates HIV-1 immature capsid assembly (26), but its exact mechanism of action has not been defined. In addition to ABCE1, the cellular DEAD box helicase 6 (DDX6) is also associated with HIV-1 assembly intermediates (27). Knockdown of DDX6 resulted in reduction of released immature capsids without affecting steady-state Gag levels, suggesting a role for DDX6 in facilitating HIV-1 capsid assembly; moreover, rescue of DDX6-depleted cells with wild-type (WT) DDX6, but not an ATPase mutant of DDX6, established a role for the RNA helicase activity of DDX6 in promoting HIV-1 immature capsid assembly (27). In keeping with our finding that ABCE1 and DDX6 are associated with Gag in immature capsid assembly intermediates and released upon completion of the immature capsid, a proteomics study found that ABCE1 and DDX6 are associated with full-length Gag in cells, but are not present in the virus (28).

Interestingly, DDX6 is a well-studied marker found in cellular RNA granules, which are host complexes involved in regulation and metabolism of RNA in the cytoplasm (reviewed in (29, 30)). Although ABCE1 was not described previously as an RNA granule protein, immunoprecipitation studies showed ABCE1 and DDX6 are associated in the absence of assembling Gag (27), suggesting that ABCE1 is also found in some DDX6-containing RNA granules. Consistent with that observation, cytoplasmic HIV-1 Gag first associates with DDX6 and ABCE1 when Gag co-opts a host complex (here termed an RNA granule because it contains the RNA granule maker DDX6), thereby forming the ~80S assembly intermediate (27). Gag then targets, along with ABCE1, DDX6 and other RNA granule proteins, to the plasma membrane (22), where Gag multimerizes. The cellular RNA granule proteins then dissociate upon completion of the immature capsid, before budding and release (26, 27). If similar RNA granules are co-opted by FIV during assembly, that would support the utility of the FIV animal model for identifying compounds that might inhibit both FIV and HIV-1 assembly by interfering with these intracellular events.

Other primate lentiviruses (e.g. HIV-2, SIVmac239, SIVagm) form similar intermediates that are associated with ABCE1 (31), suggesting that co-opting a host RNA granule at an early stage of immature capsid assembly is a conserved feature among primate lentiviruses. To date, a role for ABCE1-containing RNA granules in assembly of other retroviruses has not been shown. The structural conservation of lentiviral Gags described above led us to ask whether non-primate lentiviruses also co-opt ABCE1- and DDX6-containing host RNA granules to form immature capsid assembly intermediates. Here we showed that WT FIV Gag forms complexes in feline cells that are similar in size to the assembly intermediates formed by HIV-1. To confirm that these complexes behave like assembly intermediates we first generated assembly-defective FIV Gag mutants that correspond to known assembly-defective HIV-1 Gag mutants. We then showed that these FIV Gag mutants are arrested at the same assembly intermediates as the corresponding HIV-1 Gag mutants. Additionally, we demonstrated that FIV Gag is associated with endogenous ABCE1 and DDX6 by co-immunoprecipitation in feline cells. Lastly, we utilized an FIV Gag-ABCE1 proximity ligation assay (PLA) to demonstrate the association of assembly-competent, but not assembly-incompetent, FIV Gag with ABCE1 *in situ* using fluorescence microscopy. Based on these findings, we propose that FIV Gag co-opts RNA granules containing ABCE1 and DDX6, to form intracellular assembly intermediates, and that the assembly pathway defined previously for primate lentiviruses is conserved in a non-primate lentivirus. Given the identification of small molecules that inhibit rabies virus replication by targeting ABCE1-containing assembly intermediates (32), the conservation of such intermediates among lentiviruses has important implications for the development of new antiretroviral compounds.

## Results

To study FIV immature capsid assembly, we generated three FIV Gag-expressing constructs (Fig. 1A). To generate the first construct (termed FIV), we introduced two previously described modifications into the FIV-34TF10 proviral clone (33): specifically, we restored ORFA expression (34, 35) and replaced the native FIV promoter with the CMV promoter to allow for expression in both feline and human cells (36). The second construct is an FIV proviral clone that contains the two modifications described above as well as an inactivating mutation in the viral protease (termed FIV pro-). This was generated because preventing the protease-mediated cleavage of Gag following immature capsid assembly allows Gag to remain full-length, which makes tracking and quantification by SDS-PAGE easier (Fig. 1A; (37)). Finally, since Gag alone should be able to make virus-like particles (VLPs), we also codon-optimized FIV-34TF10A Gag and cloned it into an expression vector to make the third construct (termed FIV CO-Gag), which expresses FIV Gag in the absence of other viral proteins (Fig. 1A). To validate these three constructs, the feline astrocyte cell line G355-5 was transfected and analyzed both for steady-state expression of Gag and for VLP production by western blotting (WB) with an antibody directed against FIV CA-CTD (αFIV CA). The proviral constructs, FIV and FIV pro-, resulted in intracellular FIV Gag expression and production of either mature VLPs containing cleaved CA (~24 kDa, “p24”) or immature VLPs containing uncleaved Gag (~50 kDa, “p50”), respectively (Fig. 1B), both detected by αFIV CA. Expression of FIV CO-Gag alone resulted in high steady-state levels of Gag, as expected, and release of immature VLPs (Fig. 1B). To confirm the integrity of our infectious FIV construct, we demonstrated that virus released from G355-5 cells transfected with this construct displayed abundant reverse transcriptase activity (Fig. 1C). Moreover, when used to infect G355-5 cells, this virus stock caused a spreading infection as indicated by increased reverse transcriptase activity and syncytia formation over time (D. Ressler and J.C. Reed, unpublished observations). To confirm that the non-infectious constructs assembled VLPs properly, we used equilibrium centrifugation to determine the density of immature VLPs produced from cells transfected with these constructs. In our experiments, the average density of FIV immature VLPs (with envelopes intact) ranged from 1.12-1.16g/mL (N. Westergreen and J.C. Reed, unpublished observations), which is close to the published FIV VLP density estimate of 1.15 g/mL (Pedersen, 1987). Thus, measurements of reverse transcriptase activity and buoyant densities showed that all three of our constructs produced properly formed VLPs.

**Figure 1.**
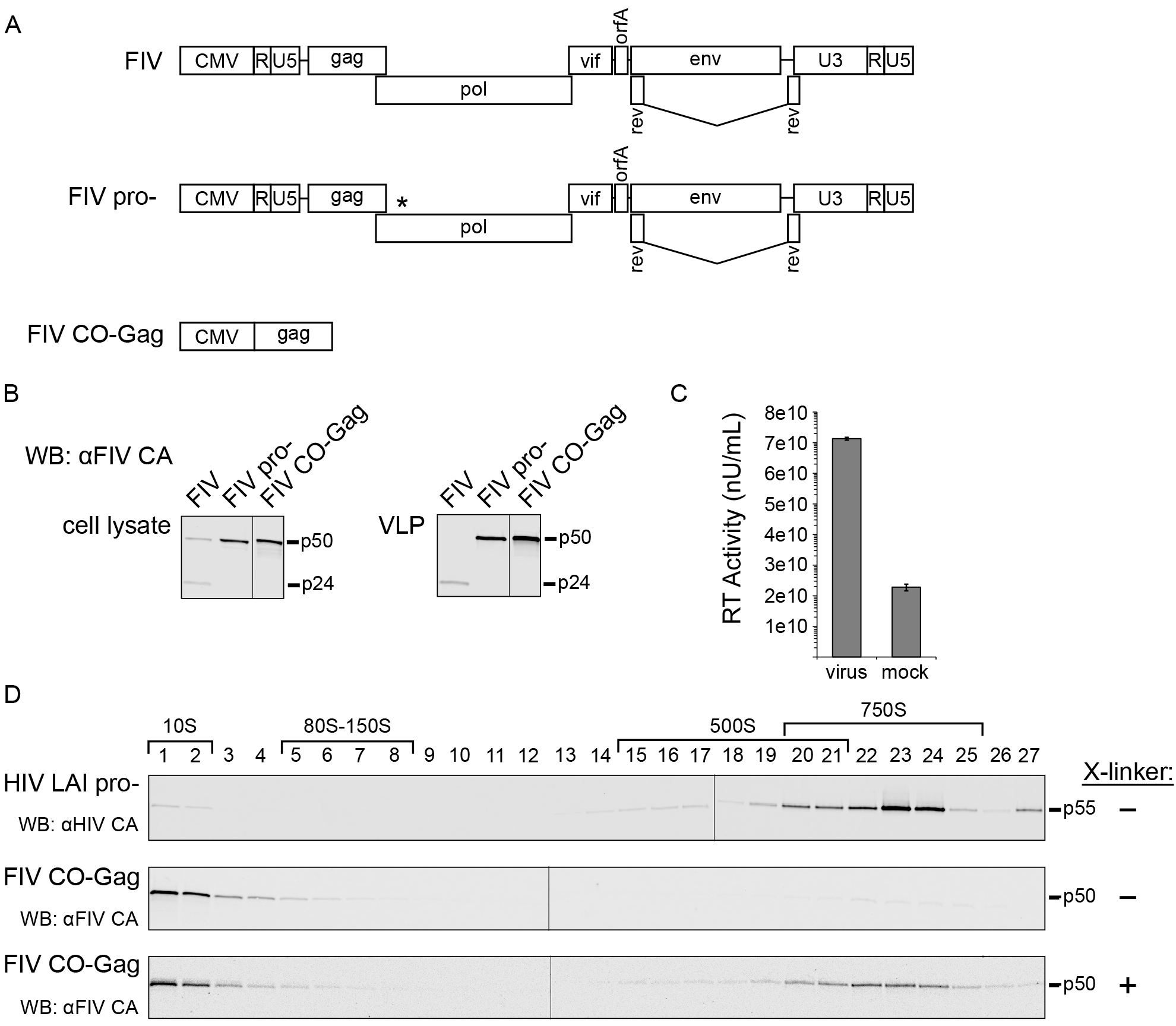
FIV immature capsids are unstable when de-enveloped, but stabilized by cross-linking. (A) Diagrams depicting gene maps of the FIV proviral construct (FIV), the FIV proviral construct containing an inactivated protease gene (FIV pro-), and codon-optimized FIV Gag alone (FIV CO-Gag). (B) Each construct was expressed in feline G355-5 cells. Cell lysates and pelleted virus-like particles (VLP) from media were analyzed by WB with an antibody to FIV CA-CTD (αFIV CA). (C) Media was collected from 293T cells transfected with FIV or mock transfected and examined for reverse transcriptase activity by SG-PERT. Standards processed in parallel were used to determine nU/mL of reverse transcriptase activity. (D) Pelleted VLPs produced either by COS-1 cells transfected with HIV-1 LAI pro- or G355-5 cells transfected with FIV CO-Gag were de-enveloped and subjected to velocity sedimentation followed by WB with an antibody to HIV-1 Gag CA-CTD (αHIV CA) or with αFIV CA. Symbols (+/−) at the right indicates whether VLPs were cross-linked (X-linked) with DSP prior to removal of the viral envelope. Approximate S-values are indicated with brackets above the WB of gradient fractions, and expected migrations of HIV-1 Gag p55 and FIV Gag p50 are shown to the right. Data shown are from a single experiment and are representative of two repeats.

Based on studies of HIV-1 assembly intermediates, we expect FIV assembly intermediates to be smaller than completed FIV immature capsids. For this reason, we first defined the migration of completed FIV immature capsids in velocity sedimentation gradients used previously to separate HIV-1 assembly intermediates (24). Immature VLPs released from cells expressing FIV CO-Gag or HIV-1 LAI pro-provirus were isolated from the culture media, de-enveloped with non-ionic detergent to release the immature capsid, and analyzed by velocity sedimentation. The HIV-1 immature capsids generated in this manner migrated at ~750S in the velocity sedimentation gradient, as described previously ((20, 24); Fig. 1D, upper panel). However, when the FIV immature capsids generated in this manner were analyzed in parallel, FIV Gag was found only in the soluble (~10S) region of the gradient (Fig 1D, middle panel). Given that VLPs of the correct density were observed when envelopes were intact, as described above, the lack of intact immature capsids following de-envelopment suggested that Gag in FIV immature capsids disassociates following removal of the viral envelope. Consistent with this hypothesis, immature capsids produced by de-envelopment of VLPs from cells expressing the FIV pro-proviral clone were also unstable (N. Westergreen and J.C. Reed, unpublished observations). To examine whether FIV immature capsids are intrinsically less stable than HIV-1 immature capsids, we used DSP to cross-link FIV immature capsids to preserve their integrity prior to removal of the viral envelope. Indeed, when velocity sedimentation was performed after cross-linking, de-enveloped FIV immature capsids migrated at ~750S (Fig. 1D, lower panel). Thus, FIV immature capsids appear to have a similar sedimentation value as HIV-1 immature capsids but are more labile, requiring cross-linking for stability after de-envelopment.

Having established the S-value of the completed FIV immature capsid, we next asked whether we could identify putative intracellular FIV assembly intermediates, which should be present transiently in small quantities and would likely have S-values similar to those of intracellular HIV-1 assembly intermediates: ~10S, ~80S, ~150S, and ~500S (21, 22, 26). Given the instability of the completed immature capsid we hypothesized that one or more of the putative FIV assembly intermediates might be unstable. To test this hypothesis, feline G355-5 cells were transfected to express FIV CO-Gag and prior to harvest were either mock treated or treated with DSP to cross-link putative FIV assembly intermediates. Harvested lysates were subjected to velocity sedimentation, and analyzed by WB for FIV Gag. The velocity sedimentation conditions used here are expected to optimally separate ~10S, ~80S, ~500S complexes; under these conditions, peaks in the ~80S and ~150S regions co-migrate and the ~500S and ~750S regions overlap. Intracellular steady-state Gag levels in cell lysates from mock treated and DSP cross-linked cells were comparable (Fig. 2A). In mock treated cell lysates, FIV Gag was found in complexes corresponding to the HIV-1 ~10S and ~80S assembly intermediates with only faint FIV Gag bands observed in the ~500S region (Fig. 2B, upper panel). In contrast, the DSP cross-linked cell lysates contained abundant FIV Gag in complexes corresponding to the HIV-1 ~10S, ~80S, and ~500S assembly intermediates (Fig 2B, lower panel). Taken together, these data suggest that FIV forms immature capsid assembly intermediates that are similar in size to previously described HIV-1 assembly intermediates, but with the larger ~500S FIV intermediate being less stable than its HIV-1 counterpart, as is the case with the released FIV immature capsid.

**Figure 2.**
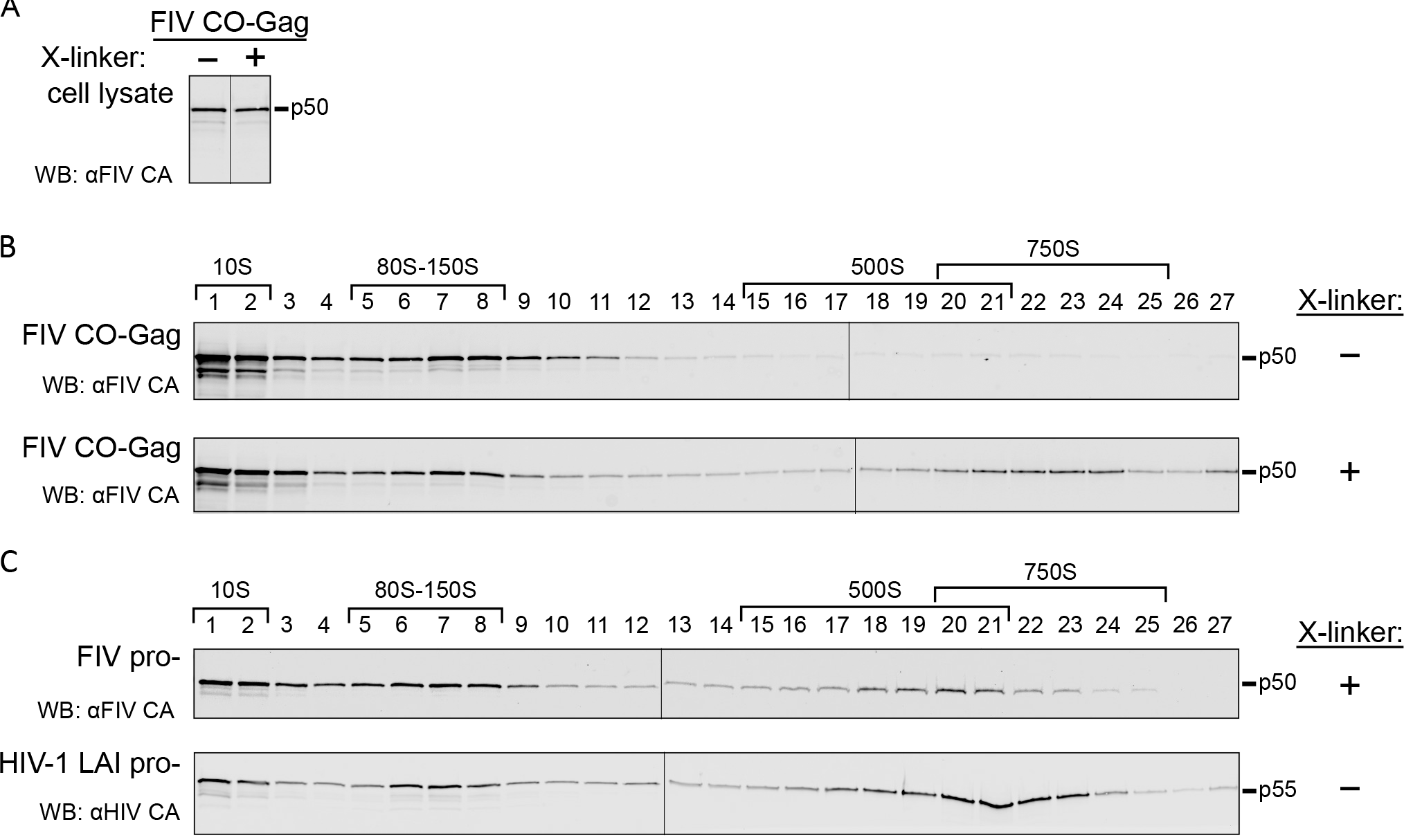
FIV produces intermediates in cells that are similar in size to HIV-1 assembly intermediates. (A) Analysis of cell lysates from G355-5 cells transfected with FIV CO-Gag that were either mock treated or treated with DSP cross-linker (X-linker) prior to harvest. WB was performed with an antibody to FIV CA-CTD (αFIV CA) and bands were taken from a single exposure of film. (B) Cell lysates blotted in panel A were subjected to velocity sedimentation in parallel, and gradient fractions were analyzed by WB with αFIV CA. (C) Cell lysates from G355-5 cells transfected with FIV pro- that were cross-linked prior to harvest or COS-1 cells transfected with HIV-1 LAI pro-were subjected to velocity sedimentation. Gradient fractions were analyzed by WB with αFIV CA or an antibody to HIV Gag CA-CTD (αHIV CA). Symbols (+/−) at right indicate whether VLPs were cross-linked with DSP prior to removal of the viral envelope. The approximate S-values of FIV Gag complexes and HIV-1 assembly intermediates are indicated with brackets above the WB, and expected migrations of HIV-1 Gag p55 and FIV Gag p50 are shown to the right.

To determine if putative FIV assembly intermediates could also be detected in cells transfected to express an FIV proviral clone, we analyzed cross-linked G355-5 cells transfected to express FIV pro-, using velocity sedimentation and WB of gradient fractions for FIV Gag (Fig 2C, upper panel). As with FIV CO-Gag, the ~10S, ~80S, and ~500S putative FIV assembly intermediates were detected. In the case of FIV pro-, the putative ~10S and ~80S intermediates and the ~500S completed immature capsids were abundant in the cross-linked lysates, with more of the late ~500S putative assembly intermediate and less of the completed ~750S capsid relative to FIV CO-Gag (Fig. 2B, lower panel). The difference in the amount of intracellular ~750S completed immature capsids present at steady state could be explained by budding kinetics that may differ with constructs and cell types, as has been observed for HIV-1 (22, 25, 27). As a positive control, COS-1 cells transfected to express HIV-1 LAI pro- were also analyzed by velocity sedimentation without prior cross-linking. WB of gradient fractions for HIV-1 Gag revealed the ~10S, ~80S, and ~500S HIV-1 assembly intermediates, as expected (Fig. 2C, lower panel). Moreover, comparison of results for HIV-1 expressing lysates confirms that FIV Gag forms complexes that are similar in size to previously defined HIV-1 assembly intermediates, with the late ~500S FIV complex being less stable than the corresponding HIV-1 late assembly intermediate. Subsequent analyses of intracellular FIV assembly intermediates were all performed following cross-linking *in situ*.

Previously, rigorous pulse-chase studies demonstrated that HIV-1 Gag complexes identified in cell lysates are part of a pathway of sequentially formed assembly intermediates; moreover, HIV-1 Gag mutants that are assembly-defective are arrested at different steps of the assembly pathway and accumulate only those assembly intermediates that precede the point of arrest, further demonstrating the sequential progression of Gag through the pathway (Fig 3A;(21, 22, 24, 25)). Thus, to further test the hypothesis that the FIV Gag-containing complexes that we identified in cells (Fig. 2B) are assembly intermediates, we generated five FIV Gag mutants that are predicted to be either budding-defective or assembly-defective, and determined the pattern of FIV Gag complexes produced in cells transfected with each of these mutants. If key assembly-defective FIV Gag mutants are arrested at the same complex as the corresponding assembly-defective HIV-1 Gag mutant and forms only the FIV Gag-containing complexes that precede the point of arrest, then FIV Gag complexes are likely part of a pathway of sequential assembly intermediates analogous to the HIV-1 assembly pathway.

**Figure 3.**
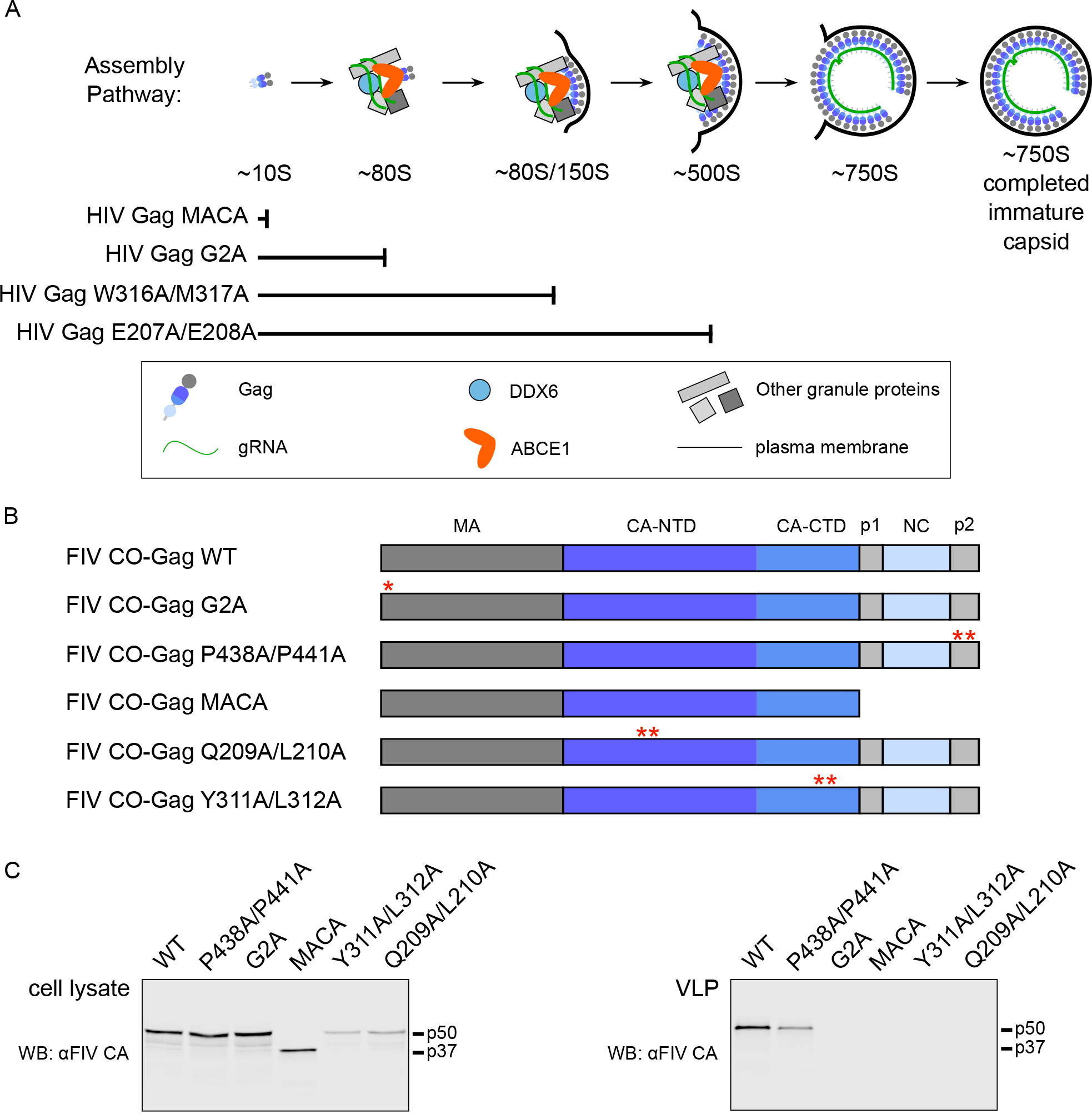
Defining VLP phenotypes of FIV Gag mutants. (A) HIV-1 assembly pathway showing the step at which specific HIV-1 Gag mutants are known to be arrested (22). The legend (in box) shows colors and symbols representing HIV-1 Gag, genomic RNA (gRNA), DDX6, ABCE1, other granule proteins, and the plasma membrane. Point mutations are numbered relative to the start of HIV-1 Gag. (B) Diagram of WT FIV Gag and FIV Gag mutants used in this study, with point mutations indicated by red asterisks. MA, dark grey; CA-CTD, dark blue; CA-NTD, blue; NC, light blue; spacer peptide (p1) and late domain (p2), light grey. FIV viral protease cleavage sites are diagramed as vertical black lines and the names of Gag domains are indicated above the diagrams. Mutated residues are numbered relative to the start of Gag. (C) Equivalent aliquots of cell lysates and pelleted virus-like particles (VLP) from G355-5 cells transfected with WT FIV CO-Gag or the indicated FIV CO-Gag mutants were analyzed by WB with an antibody to FIV CA-CTD (αFIV CA). Expected migrations for full-length (p50) and MACA (p37) Gag proteins are shown to the right. Data shown are from a single experiment and is representative of two repeats.

To generate a budding-defective mutant (FIV CO-Gag P438A/P441A), we altered the TSG101 binding motif in FIV p2 from PSAP to ASAA, which is known to reduce production of FIV VLPs (Fig. 3B; (11)). Given that the analogous PTAP mutation in HIV-1 inhibits virus budding but not Gag multimerization (38, 39), we would expect FIV CO-Gag P438A/P441A to be assembly-competent and therefore form all the assembly intermediates, but subsequently fail to complete budding and release. We also generated two FIV Gag mutants that are expected to be assembly-defective. One is truncated at the end of CA, resulting in expression of only MA-CA (FIV CO-Gag MACA; Fig. 3B). We would expect that, like HIV-1 MACA (Fig. 3A;(22)), FIV MA-CA would fail to produce VLPs, would be arrested as the ~10S assembly intermediate, and would thereby mark the first step in the assembly pathway.

Moreover, a similar truncation mutant in FIV Gag, FIV MA-CA-p1, also does not produce VLPs (9). In the second FIV Gag mutant, the glycine at position 2 of FIV Gag was substituted with an alanine to generate FIV CO-Gag G2A (Fig. 3B); this mutation is known to reduce VLP production (8). The analogous HIV-1 Gag G2A mutation fails to undergo myristoylation and subsequent targeting to the plasma membrane (PM) site of assembly, and is arrested as a cytosolic ~80S assembly intermediate (Fig. 3B; (22)). Hence, we expected FIV CO-Gag G2A to be arrested at the ~80S assembly intermediate. To verify that these three constructs express in cells and display the expected phenotype in assembly or budding, we examined steady-state intracellular Gag levels as well as VLP production (Fig. 3C). The budding mutant, FIV Gag P438A/P441A, was expressed at a level comparable to WT FIV CO-Gag and displayed a three-fold reduction in VLP release relative to WT (Fig. 3C). Although this budding defect was not as profound as observed previously (11), this may reflect differences in cell lines (40). For the two other FIV Gag mutations described thus far (G2A and MACA), almost no VLPs were released from transfected cells, despite intracellular Gag levels similar to WT FIV Gag (Fig. 3C), confirming that both are assembly-defective.

Having established that these three FIV Gag mutants (P438A/P441A, G2A, and MACA) display the expected VLP phenotypes, we expressed them in cells at similar steady-state levels (Fig. 4A) and asked whether they are arrested in the assembly pathway in a manner similar to the comparable HIV-1 Gag mutants (Fig. 4B and 4C). Velocity sedimentation analysis of cell lysates revealed that FIV CO-Gag produces the expected pattern of putative intracellular assembly intermediates, displaying a prominent ~80S complex and a less prominent ~500S complex as well as the ~750S completed immature capsid. As expected, the budding-defective P438A/P441A mutant produced all the putative assembly intermediates formed by FIV CO-Gag, consistent with immature capsid assembly being unaffected by mutation of the TSG101 binding motif that is important for budding but not assembly. Interestingly, the budding-defective P438A/P441A mutant accumulated more ~750S completed immature capsids than WT FIV CO-Gag, consistent with the completed P438A/P441A immature capsids being arrested at the PM before undergoing release. In contrast, FIV CO-Gag MACA only produced the ~10S complex, and FIV CO-Gag G2A produced only the ~10S and ~80S complexes (Fig. 4B and 4C). Thus, the assembly-incompetent FIV CO-Gag MACA mutant and assembly-defective FIV CO-Gag G2A mutant were each arrested at the same Gag-containing complex as their HIV-1 counterparts (Fig. 3A; (22)). Taken together, these data further support the model that intracellular FIV Gag assembles via a stepwise pathway of assembly intermediates, analogous to those described previously for HIV-1, and that these assembly intermediates can be identified at steady state in cell lysates.

**Figure 4.**
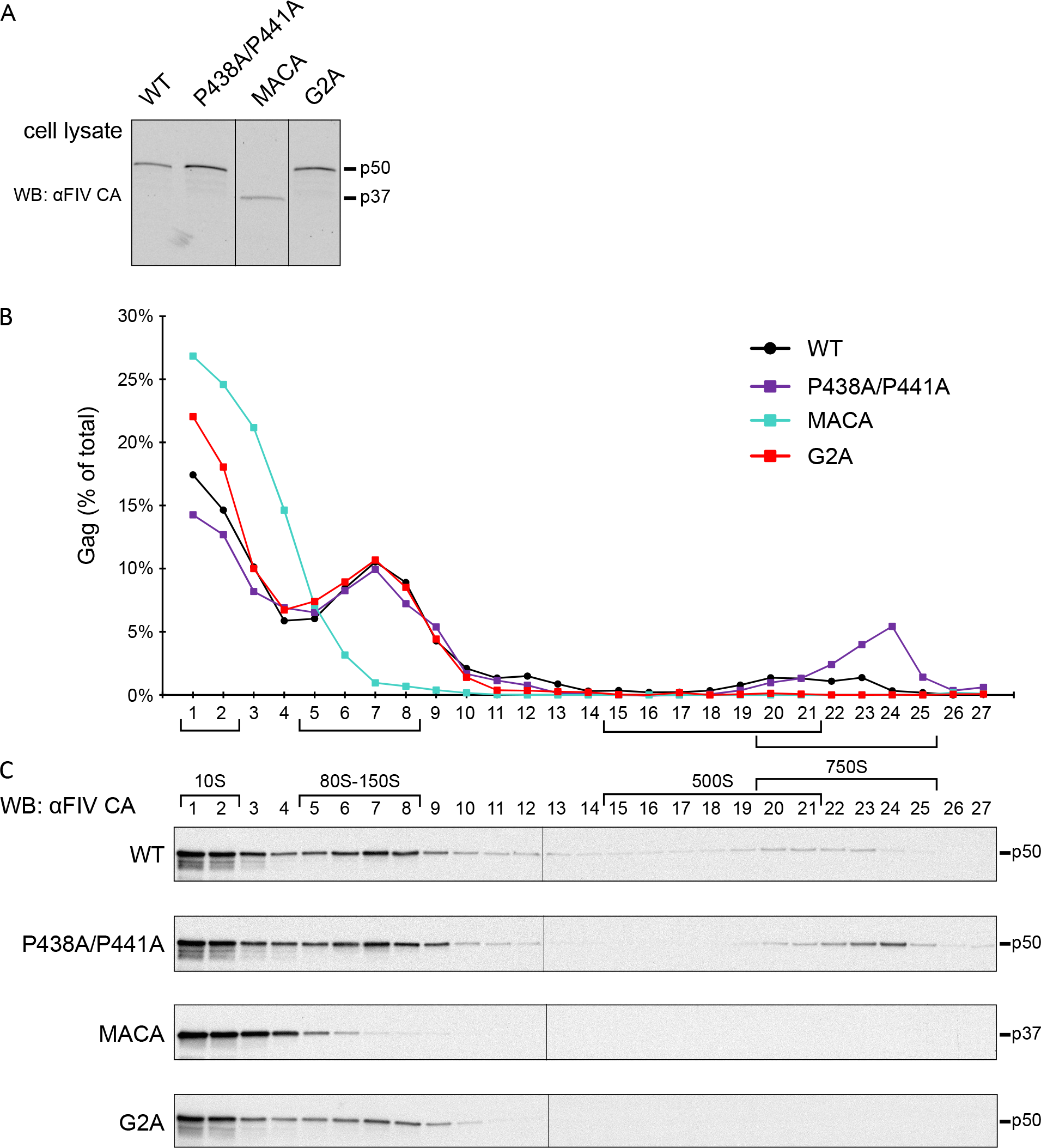
Assembly-defective FIV Gag MACA and G2A are arrested at putative assembly intermediates. (A) Equivalent aliquots of cell lysate from G355-5 cells transfected with FIV CO-Gag WT or indicated FIV CO-Gag mutants were analyzed for steady-state FIV Gag expression by WB with an antibody to FIV CA-CTD (αFIV CA). (B, C) Lysates shown in panel A above were also subjected to velocity sedimentation and gradient fractions analyzed by WB with αFIV CA. Graph displays quantification of blots shown in panel C, with the amount of Gag in each fraction shown as a percentage of total Gag in the gradient. The approximate S-values of FIV Gag-containing complexes are indicated with brackets below the graph and above the gradient WB, and expected migrations for full-length (p50) and MACA (p37) Gag proteins are shown to the right. Data shown are from a single experiment and are representative of two repeats.

Given the important role of the CA domain in HIV-1 immature capsid assembly, we also wanted to identify point mutations in FIV CA that result in assembly defects and determine if they also cause arrest of FIV Gag in the putative assembly pathway. While point mutations in CA-NTD and-CTD that impair HIV-1 immature capsid assembly have been defined ((16, 24, 41–45)), they have not yet been identified for FIV. Such FIV point mutants would allow a finer mapping FIV CA function and would allow us to compare arrest of FIV and HIV assembly-defective mutants, thereby providing additional insights into assembly mechanisms. Since high-resolution structures of the HIV-1 CA-NTD and CA-CTD are available, we sought to identify FIV residues that correspond to known HIV-1 assembly-critical residues by aligning the structure of the FIV CA subdomains onto the structure of the HIV-1 CA subdomains (Fig. 5A). For CA-CTD, we aligned the published FIV CA-CTD structure (PDB accession number 5DCK; (18)) with the HIV-1 CA-CTD domains from a published structure of HIV-1 CA (PDB accession number 5L93; (46)). However, because a high-resolution structure of FIV CA-NTD has not been reported, we generated a model of the FIV CA-NTD structure, using SWISS-MODEL, with the published RELIK CA-NTD structure as a template (PDB accession number 2XGU; (19)). This FIV CA-NTD model was then aligned with the published HIV-1 CA-NTD structure (PDB accession number 5L93; (46); Fig. 5A).

**Figure 5.**
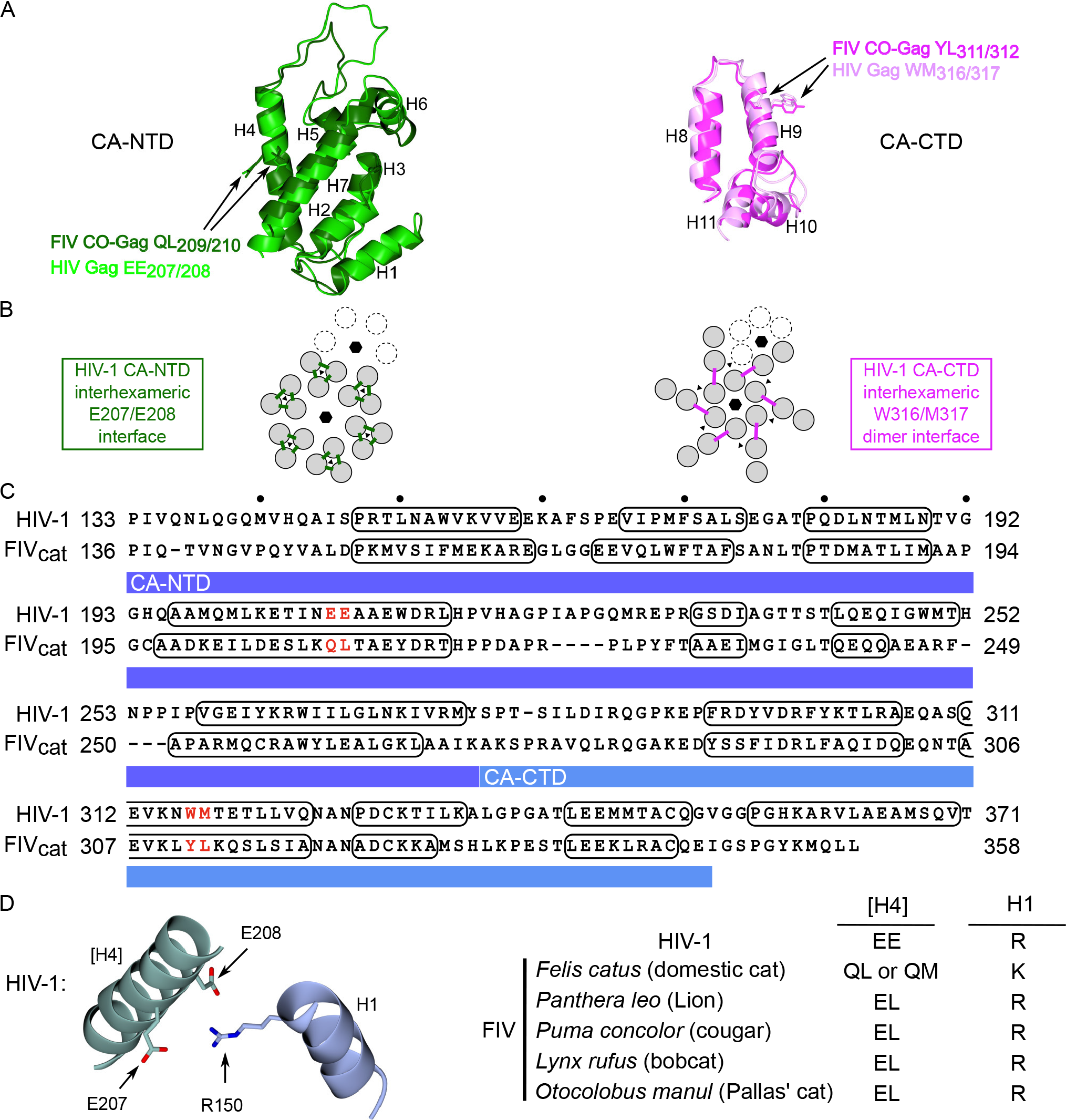
Structural homology modeling allows identification of candidate CA residues critical for FIV capsid assembly. (A) Superposition of HIV-1 CA-NTD (light green; PDB accession number 5L93) and a homology modeled structure of FIV CA-NTD (dark green) using RELIK CA-NTD as a template (PDB accession number 2XGU) at left; and HIV-1 CA-CTD (light pink; PDB accession number 5L93) and FIV CA-CTD (dark pink; PDB accession number 5DCK) at right. Side chains of FIV residues selected for mutation, and their HIV-1 counterparts, are shown in the same color as the main chain. CA alpha helices 1-11 are numbered (H1-H11). (B) Diagram of the HIV-1 immature capsid structure depicting the six-fold and three-fold points of symmetry in the CA-NTD layer (black hexagons and black triangles, respectively; at left) and CA-CTD layer (at right), which were previously identified in a high-resolution structure of the completed immature capsid (46). Dotted subunits indicate an adjacent hexamer. The HIV-1 CA-NTD inter-hexameric interface studied here is shown by green lines, but is shown only for the central hexamer. The HIV-1 CA-CTD inter-hexameric interface studied here is shown by pink lines, but is shown only for the central hexamer. (C) Sequence alignment is based on a structural alignment of CA-NTD and CA-CTD from sources described in section A above. Numbering at right and left of each line is from the start of Gag. Residues forming alpha-helical structures are outlined with black ovals. CA-NTD and CA-CTD are highlighted with a dark blue and light blue bar respectively. Red letters indicate residues chosen for mutation (Q209A/L210A and Y311A/L312A). (D) Diagram of the residues proposed to form a salt bridge in the HIV-1 immature capsid structure (R150 in helix 1 (H1) with E207/E208 in helix 4 ([H4]); brackets indicate helix in neighboring 3-fold symmetry mate; adapted from (46)). Table lists consensus resides at these positions in helix 4 and helix 1 for HIV-1, FIV *Felis catus* (domestic cat), FIV *Panthera leo* (lion), FIV *Puma concolor* (cougar), FIV *Lynx rufus* (bobcat), FIV *Otocolobus manul* (Pallas’ cat).

After aligning the FIV CA subdomain structures on an HIV-1 CA structure (Fig. 5A), we next asked which residues in the FIV structures most closely correspond to assembly-critical HIV CA residues. In CA-NTD, residues in helix 4 were of particular interest because helix 4 forms an exposed surface ((44)) and is critical for inter-hexameric CA-NTD contacts in a high-resolution cryoEM structure ((46); Fig. 5B). In keeping with this, the HIV-1 Gag helix 4 mutant E207A/E208A (Fig. 5C) displays reduced VLP production (44) but forms all the assembly intermediates, including the late ~500S assembly intermediate (22), indicating arrest just prior to formation of the ~750S completed capsid (Fig. 3A). Moreover, a recent cryoEM structure of HIV-1 capsid suggests that E207 and E208 in CA-NTD helix 4 may participate in a salt bridge with R150 in CA-NTD helix 1 of a neighboring 3-fold symmetry mate ((46); Fig. 5D). This interaction may contribute to stabilizing the immature HIV-1 capsid; thus, it is possible that the observed arrest of the E207/E208 mutant in the assembly pathway is explained by disruption of this salt bridge. CA-NTD helix 4 is also important for FIV immature capsid assembly since its deletion reduces VLP production (9). Thus, we chose to mutate the FIV residues Q209/L210 to alanine based on their alignment with E207/E208 in the superimposed HIV-1 and FIV CA-NTDs structure (Fig. 5A and C). Additionally, the side chains of Q209/L210 residues had a similar orientation relative to the corresponding E207/E208 residues in the HIV-1 CA structure (Fig. 5A), further supporting the selection of Q209/L210 for mutation. Consistent with their similar alignment and orientation, QL (or QM) residues are highly conserved at this position in FIV variants from domestic cats (present in 312 out of 322 FIV Gag domestic cat sequences deposited to NCBI; Fig. 5D and Suppl. Table 1). Notably, as described in the Discussion section, the finding that Q209 in FIV CA-NTD of domestic cats corresponds to E209 in HIV-1 CA-NTD based on alignment and orientation raises some interesting structural issues, since unlike glutamic acid, glutamine would not be expected to form a salt bridge in FIV Gag due to its lack of a negative charge.

To test whether the FIV Q209/L210 residues are analogous to the assembly-critical HIV-1 E207/E208 residues, we introduced Q209A/L210A mutation into FIV CO-Gag and examined the effect on VLP production and progression through the assembly pathway. We found that VLP production by FIV CO-Gag Q209A/L210A was dramatically reduced despite intracellular Gag expression (Fig. 3C), thus recapitulating the VLP defect observed for the corresponding HIV-1 mutant (22, 44). When expressed in cells at similar intracellular levels as WT FIV CO-Gag (Fig. 6A), FIV CO-Gag Q209A/L210A formed the ~10S, ~80S, and ~500S complexes, but little to no ~750S completed immature capsid, as indicated by velocity sedimentation analysis (Fig. 6B and C). From these data, we conclude that FIV CO-Gag Q209A/L210A is arrested at the ~500S assembly intermediate, since it forms the ~500S intermediate (Fig. 6C) but fails to complete assembly and release (Fig. 3C), as observed for the corresponding HIV-1 E207A/E208A mutant (Fig. 3A; (22)).

**Figure 6.**
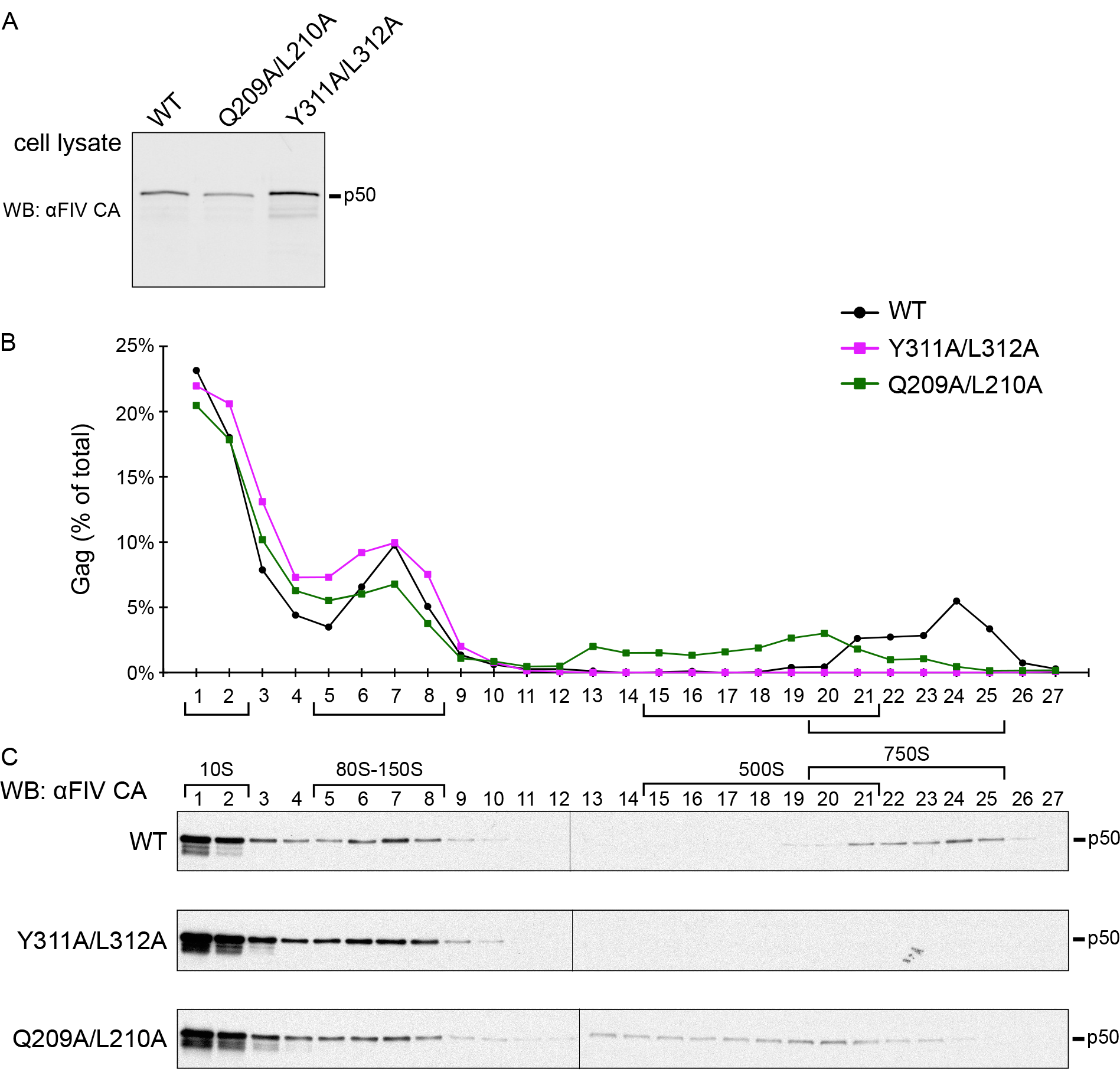
Assembly-defective FIV Gag CA-NTD and CA-CTD mutants are arrested at putative assembly intermediates. (A) Equivalent aliquots of cell lysate from G355-5 cells transfected with FIV CO-Gag WT or indicated FIV CO-Gag mutants were analyzed for steady-state FIV Gag expression by WB with an antibody to FIV CA-CTD (αFIV CA). (B, C) Lysates shown in panel A above were also subjected to velocity sedimentation and gradient fractions analyzed by WB with αFIV CA. Graph displays quantification of blots shown in panel C, with the amount of Gag in each fraction shown as a percentage of total Gag in the gradient. The approximate S-values of FIV Gag-containing complexes are indicated with brackets below the graph and above the gradient WB, and expected migration for full length (p50) Gag protein are shown to the right. Data shown are from a single experiment and are representative of two repeats.

Similarly, we also sought to identify residues in FIV CA-CTD that correspond to assembly-critical HIV-1 residues. In HIV-1 CA-CTD, CA helix 9 mediates an important inter-hexameric CA-CTD dimerization interface in the completed immature capsid ((16); Fig. 5B). We would expect that FIV helix 9 also forms this critical CA-CTD dimer interface, especially given that FIV CA-CTD can functionally replace SIV CA-CTD, as shown with SIV-FIV chimeras (47). It was demonstrated previously that the W316/M317 mutation within helix 9 of HIV-1 CA inhibits VLP production (22, 44) and arrests HIV-1 Gag assembly at the ~80S intermediate (Fig 3A; (22)). To determine if FIV assembly also depends on this critical CA-CTD dimerization interface, FIV Gag residues Y311 and L312 were chosen for mutation based on their alignment with, and similar orientation to, residues W316 and M317 in HIV-1 helix 9 (Fig. 5A and C).

When expressed in cells, FIV CO-Gag Y311A/L312A displayed dramatically reduced VLP production (Fig. 3C), thus recapitulating the VLP defect observed for the analogous HIV-1 Gag W316/M317 (22, 44). Velocity sedimentation analysis revealed that FIV CO-Gag Y311A/L312A produced only the ~10S and ~80S complexes, while WT CO-Gag produced 10S, ~80S, and ~500S/750S intermediates when expressed at similar steady-state intracellular levels and analyzed in parallel (Fig. 6A-C). Thus, FIV CO-Gag Y311A/L312A is arrested at the ~80S putative assembly intermediate as observed for the corresponding HIV-1 W316A/M317A mutation (Fig. 3A; (22)). Together, analysis by velocity sedimentation shows that each of the four assembly-defective Gag mutants that we examined is arrested at the same assembly intermediate as its HIV-1 counterpart (Figs. 4 and 6; summarized in Fig. 7A).

**Figure 7.**
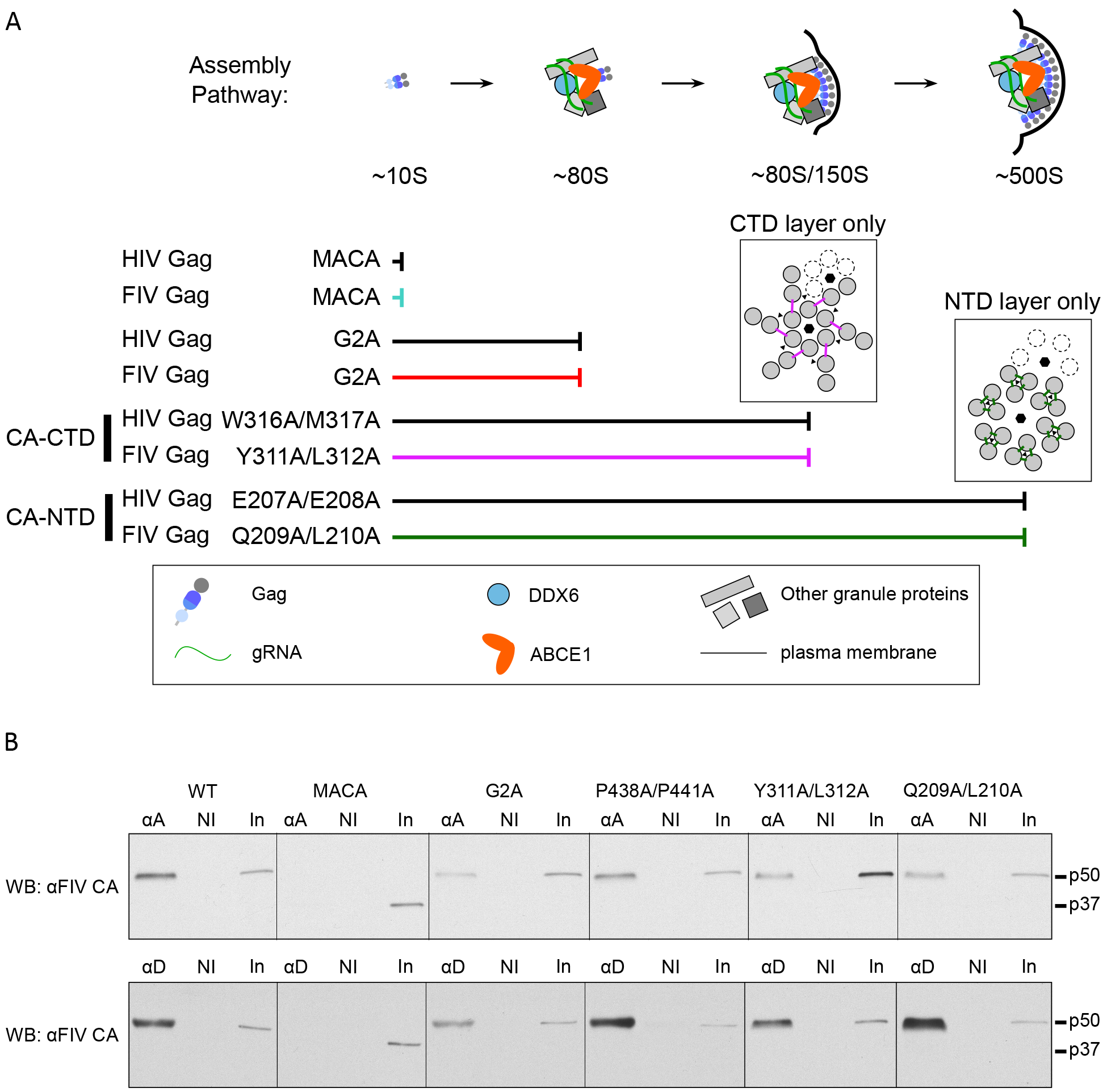
FIV Gag, like HIV-1 Gag, is associated with both ABCE1 and DDX6 by co-immunoprecipitation. (A) FIV assembly pathway model showing that FIV Gag mutants are arrested in the assembly pathway at the same points as their HIV-1 counterparts. Located above the point of arrest of the CA-CTD mutations and the CA-NTD mutations are diagrams of the HIV-1 immature capsid structure showing either the HIV-1 CA-CTD layer only (left) or the HIV-1 CA-NTD layer only (right). The six-fold and three-fold points of symmetry are indicated (black hexagons and black triangles, respectively) and the dotted subunits indicate an adjacent hexamer. The HIV-1 CA-NTD inter-hexameric interface studied here is shown by green lines, but is shown only for the central hexamer. The HIV-1 CA-CTD inter-hexameric interface studied here is shown by pink lines, but is shown only for the central hexamer. These diagrams are based on the high-resolution structure of the immature capsid described in (46). The legend (in box) shows colors and symbols representing Gag (FIV or HIV-1), genomic RNA (gRNA), DDX6, ABCE1, other granule proteins, and the plasma membrane. (B) Lysates of G355-5 cells transfected with the indicated constructs were subjected to immunoprecipitation with an antibody to ABCE1 (αA) or an antibody to DDX6 (αD) alongside a non-immune (NI) control. Immunoprecipitation eluates and an aliquot of input (In) were analyzed by WB with an antibody to FIV CA-CTD (αFIV CA).

To further confirm that the FIV Gag-containing complexes are assembly intermediates, we asked if they contain the same cellular proteins found in HIV-1 assembly intermediates. The ~80S and ~500S HIV-1 capsid assembly intermediates contain several host proteins, of which the best studied are the enzymes ABCE1 (26) and DDX6 (27). Both ABCE1 and DDX6 facilitate events in HIV-1 assembly (26, 27) and serve as markers for assembly intermediates. Thus, we asked whether FIV Gag is also associated with endogenous ABCE1 and DDX6 in feline G355-5 cells. FIV CO-Gag was expressed in G355-5 cells and cell lysates were subjected to immunoprecipitation with either antibody to human ABCE1 or antibody to human DDX6 alongside a non-immune control, followed by WB with an antibody to FIV CA-CTD to detect co-immunoprecipitated FIV Gag. Because both ABCE1 and DDX6 are highly conserved between humans and domestic cat (99.8% identity) and the ABCE1 and DDX6 antibodies we used (described previously (26, 27)) are directed against peptides with 100% identity between humans and domestic cat, we expected both antibodies to recognize the feline homologs. Indeed, we found that WT FIV Gag was associated with endogenous feline ABCE1 and DDX6 by immunoprecipitation (Fig. 7B). Thus, like HIV-1 Gag, FIV Gag appears to co-opt an ABCE1- and DDX6-containing RNA granule.

The ~10S HIV-1 assembly intermediate, which corresponds to soluble Gag, is not associated with ABCE1 and DDX6 (26, 27, 31), indicating that HIV-1 Gag likely associates with ABCE1 and DDX6 after it targets to an RNA granule. Given this, we would not expect the assembly-incompetent FIV MACA, which is arrested at the ~10S assembly intermediate (Fig. 4C), to be associated with ABCE1 and DDX6. In contrast, we would expect all the other assembly-defective mutants studied here (FIV G2A, Q209A/L210A and Y311A/L312A) to associate with ABCE1 and DDX6, since they are all arrested at either the ~80S intermediate (G2A and Y311A/L312A; Fig. 4C and 6C) or the ~500S intermediate (Q209A/L210A; Fig. 6C). Additionally, the assembly-competent P438A/P441A mutation, which forms all the assembly intermediates (Fig. 4C), should also associate with ABCE1 and DDX6. Indeed, as expected, all the FIV Gag mutants we studied here, with the exception of FIV MACA, were associated with endogenous ABCE1 and DDX6 in feline G355-5 cells by co-immunoprecipitation (Fig. 7B). These findings support a model in which FIV Gag localizes to an RNA granule that contains ABCE1 and DDX6, as shown previously for HIV-1.

Finally, to validate the association of FIV Gag with ABCE1 we asked whether ABCE1 is co-localized with Gag in intact cells. For this purpose, we utilized the proximity ligation assay (PLA), which will only produce fluorescent spots at sites where two proteins are within 40 nm of each other *in situ*. Briefly, proteins of interest (FIV Gag and ABCE1) are labeled with primary antibodies, which in turn are detected by a pair of secondary antibodies that are each conjugated to one of two complementary oligonucleotides; the two secondary antibodies are ligated via annealing between the conjugated complementary oligonucleotides only when the proteins of interest are within 40 nm of one another. Reagents added subsequently result in a rolling circle amplification product that is recognized by a fluorophore-conjugated oligonucleotide, creating a punctate fluorescent signal wherever co-localization occurs (Fig. 8A; (48)). Thus, if FIV Gag is in close proximity with ABCE1 within an ABCE1-containing RNA granule, cells expressing WT FIV Gag would be expected to display abundant Gag-ABCE1 PLA spots. In contrast, cells expressing the assembly-defective FIV Gag MACA mutant would be expected to display few PLA spots, since FIV Gag MACA is arrested at the ~10S intermediate and does not associate with ABCE1 by coIP. To avoid FIV-Env induced cytopathicity (36) that might interfere with the PLA assay, we utilized proviral constructs that only expressed a truncated form of Env (FIV pro- env-, diagrammed in Fig. 8A). We found that cells transfected with FIV pro-env-encoding WT Gag (WT, Fig. 8A) contained an average of 34.9 Gag-ABCE1 PLA spots per cell (Fig. 8B, red spots in Fig. 8C second column of top row), and concurrent FIV Gag indirect immunofluorescence staining showed that majority of these PLA spots occurred in cells expressing FIV Gag at low or high levels (Fig. 8C, first and third column of top row). In contrast, cells transfected with FIV pro- env- encoding the assembly-defective FIV Gag MACA mutant (MACA, Fig. 8A) contained nearly four-fold fewer Gag-ABCE1 PLA spots per cell than WT FIV Gag (an average of 9.5 Gag-ABCE1 PLA spots per cell; Fig. 8B, red spots in Fig. 8C second column of bottom row), despite abundant FIV MACA staining by indirect immunofluorescence (Fig. 8C, first and third column of bottom row). Note that because FIV WT and mutant Gag and ABCE1 are both present in the ~10S region of velocity sedimentation gradients (Fig. 4C and 6C; J.C. Reed and J.R. Lingappa, unpublished observations), these proteins may be in proximity to some extent in the soluble fraction of the cytoplasm, outside of RNA granules. This could explain the presence of a background level of PLA spots for FIV MACA. These data show that FIV Gag is associated with ABCE1 *in situ*; moreover, together with the co-immunoprecipitation data, the PLA data further support a model in which assembling FIV Gag co-opts a DDX6 and ABCE1-containing RNA granule.

**Figure 8.**
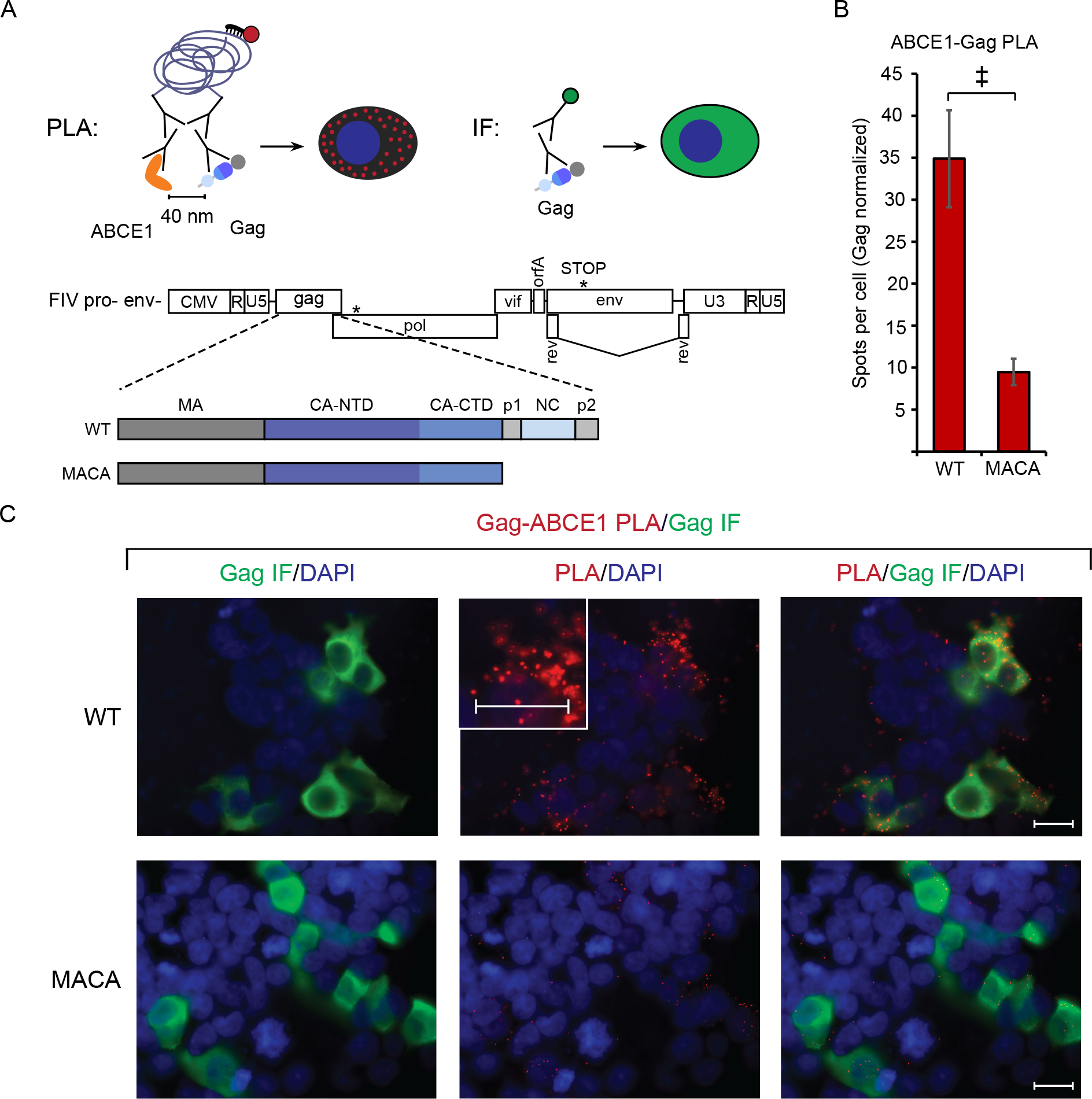
PLA confirms co-localization of FIV Gag and ABCE1 *in situ* upon provirus expression. PLA spots indicate regions in which Gag is within 40 nm from ABCE1 *in situ*, with concurrent αGag immunofluorescence (IF) allowing quantification of intracellular Gag levels. (A) Experimental schematic showing fluorophore detection of the rolling-circle amplification product from ligated oligonucleotides attached to mouse and rabbit secondary antibodies. These detect primary antibodies directed against FIV Gag and ABCE1. Also shown is the gene map of the FIV pro- env- proviral construct, with the point mutation in protease and premature stop codon in env marked by asterisks. PLA experiments utilized the FIV pro-env-provirus encoding either WT Gag or MACA, diagrammed below the gene map. (B) Following transfection of these proviruses into 293T cells and processing for PLA, the average number of PLA spots per cell was determined for all Gag-IF-positive cells in five randomly chosen fields, and normalized to Gag levels as measured by IF signal intensity. Error bars indicate SEM. Double cross indicates a significant difference relative to WT (p<0.01 in a two-tailed, paired t-test). (C) Panels show representative images from each transfection. Columns from left to right for each construct: Gag IF (green) with DAPI-stained nuclei (blue), FIV Gag-ABCE1 PLA signal (red) with DAPI-stained nuclei (blue), and a merge of images in the first two columns. The center PLA panel in the top row contains an inset showing a high magnification view of the cell to the right of the inset. Scale bars, 5 μM.

## Discussion

Here we present evidence that, in feline cells, the non-primate lentivirus FIV Gag assembles immature capsids via a pathway of RNA-granule-derived assembly intermediates, as shown for HIV-1 and other primate lentiviruses (20–22, 26, 27, 31). Support for this comes from four findings: first, FIV Gag forms complexes in cells that resemble well-studied HIV-1 immature capsid assembly intermediates in size and shape (Fig. 2); second, assembly-defective FIV Gag mutants are arrested in the putative assembly pathway and the points of arrest are the same as for the analogous HIV-1 Gag assembly-defective mutants (Fig. 4 and 6); third, FIV Gag is associated with the RNA granule proteins ABCE1 and DDX6 by co-immunoprecipitation (Fig. 7B), as observed for HIV-1 Gag in assembly intermediates; and fourth, assembling FIV Gag colocalizes with ABCE1 *in situ*, while an assembly-defective mutant does not (Fig. 8). Taken together these data argue that the ~80S and ~500S FIV Gag-containing complexes correspond to early and late immature capsid assembly intermediates, respectively, and that the immature assembly pathway defined by studies of HIV-1 is conserved between primate and non-primate lentiviruses (Fig. 7A). Below we will use the term assemblysome to refer to collectively RNA-granule derived capsid assembly intermediates of different sizes (e.g. the ~80S and ~500S HIV-1 and FIV assembly intermediates).

Our finding that both primate (HIV-1, HIV-2, SIV) and non-primate (FIV) lentiviruses appear to form assemblysomes by co-opting the same subclass of ABCE1- and DDX6-containing RNA granules suggests that many or all Gag proteins have evolved to utilize this cellular complex. While many questions about these RNA granules remain unanswered, this conservation argues that these granules play an important role in the retroviral lifecycle. Other types of RNA granules, such as P bodies and stress granules, transiently store non-translating cellular mRNA (reviewed in (49)); thus, it is possible that these ABCE1- and DDX6-containing complexes could also be sites for storing non-translating cellular mRNA. Moreover, because FIV and HIV-1 unspliced RNA resembles cellular mRNA in that it is both capped and polyadenylated, one might expect unspliced FIV and HIV-1 viral RNA to also be stored in these ABCE1- and DDX6-containg granules. Consistent with this possibility, unspliced HIV-1 RNA is associated with ABCE1 in these RNA granules (22). Thus, targeting of retroviral Gag proteins to this RNA granule could be advantageous for a number of reasons. First, assembling Gag could take advantage of cellular facilitators of immature capsid assembly present in RNA granules, such as ABCE1 and DDX6 with HIV-1 (22, 25–27). Second, these RNA granules could provide a site in which Gag becomes concentrated, thereby facilitating Gag-Gag interactions. Third, RNA granule localization could bring assembling Gag into proximity with granule-associated unspliced retroviral RNA to facilitate packaging of the RNA genome into the assembling virus. Finally, these RNA granules may also shield retroviral RNA or other retroviral components from host innate immune sensing.

Many viruses are known to utilize RNA granules at various stages of their lifecycles (50). Canonical RNA granules include stress granules and P bodies, both of which are much larger than the ABCE1- and DDX6-containing RNA granule from which assembly intermediates are derived. For example, P bodies range from 100 – 300 nm in size (51) whereas our ~80S assembly intermediate should be approximately the same size as a ribosome (~25 nm, (52)). Others have reported that HIV-1 Gag is associated with RNA granule proteins including Staufen1 (53, 54), AGO2 (55), and MOV10 (56). However, HIV-1 Gag is not found in stress granules (54), whose formation is induced by stress; nevertheless, Gag has been shown to modulate formation of stress granules (54, 57, 58). Similarly, a recent study reported that HIV-1 Gag does not co-localize with P bodies by fluorescent microscopy (59). Consistent with that report (59), our recent PLA studies show that HIV-1 Gag is not typically associated with P bodies although it is associated with the P body protein DDX6 in much smaller granular structures (Barajas B and Lingappa JR, unpublished observations). Thus, P bodies, stress granules, and assemblysomes clearly represent RNA granules of different sizes and functions; however, more work will be needed to understand how they are related to each other and to the HIV-1 lifecycle. One hypothesis consistent with published data is that the ABCE1-and DDX6-containing granules that are co-opted by Gag are subunits of P bodies and/or stress granules that can also exist independent of P bodies and stress granules. In support of this hypothesis, the ABCE1- and DDX6-containing granules that are co-opted by Gag share at least two components with P bodies (DDX6 and AGO2 (27)), and DDX6 is also found in stress granules (reviewed in (60)). Thus, this subunit hypothesis would explain why Gag is associated with DDX6 and ABCE1 but does not co-localize with P bodies or stress granules, and why P bodies and stress granules can be impacted when Gag is expressed. While future studies will be needed to test these possibilities, the current study adds to a large body of evidence indicating that a common feature of Gag proteins is their ability to co-opt a unique ABCE1-and DDX6-containing RNA granule during immature capsid assembly.

The association of HIV-1 Gag with DDX6 and ABCE1 in assemblysomes was shown previously using co-immunoprecipitation and immunoelectron microscopy (21, 27). Here we showed that, like HIV-1 Gag, FIV Gag is associated with both of these host enzymes by co-immunoprecipitation. Here for the first time, we utilized a PLA assay that allowed us to demonstrate co-localization of FIV Gag with ABCE1 *in situ* using fluorescent microscopy. Unlike direct or indirect fluorescent microscopy, PLA results in fluorescent spots only when two proteins are in close proximity, thus allowing co-localization to be quantified even when proteins are present diffusely throughout the cytoplasm. Thus, taken together two different *in situ* approaches – PLA and immunoelectron microscopy – support the finding that assembling Gag proteins of various lentiviruses form ABCE1-containing assemblysomes, while assembly-incompetent MACA Gag mutants do not. Moreover, our PLA results suggest that PLA may be a promising approach for *in situ* confirmation of other retroviral-host protein associations.

Although our study found numerous similarities between FIV and HIV-1, we did find one significant difference, namely that the late ~500S FIV assembly intermediate and completed FIV immature capsids were relatively unstable compared to the corresponding HIV-1 complexes. Notably, our study demonstrated that *in situ* cross-linking overcomes the inherent lability of FIV complexes, allowing these complexes to be stabilized and analyzed. Thus, the cross-linking approach that we developed and validated here should allow assemblysomes for these and other retroviral capsids to be isolated and analyzed intact in the future.

The current study also advances our understanding of residues and interfaces critical for assembly of FIV immature capsids, and may have identified an example of co-evolution that could have structural implications. Previously, it has been shown that E207/E208 in helix 4 of HIV-1 CA-NTD is required for assembly (44) and for progression past the ~500S step in the assembly pathway (22), consistent with a study showing that these residues are critical for an inter-hexameric interface formed between helix 4 and helix 1 of a neighboring 3-fold symmetry mate (46). Here we showed that Q209/L210 in FIV CA-NTD helix 4 function in a manner analogous to E207/208 in helix 4 of HIV-1 CA-NTD, in that Q209/L210 are important for VLP production and for progression past the ~500S Gag-containing complex (Fig 6B and C). However, we also noted a difference in the HIV-1 and FIV residues that refines our understanding of this interface. Others have proposed that a salt bridge may form between the negatively charged E207 and E208 in helix 4 of HIV-1 CA-NTD and the positively charged R150 in helix 1 of HIV-1 Gag CA-NTD (E-R salt bridge; (46)); but this salt bridge is unlikely to form between the comparable FIV Gag residues, Q209/L210 (neither of which is negatively charged), and K152, which is analogous to HIV-1 R150. Our finding that FIV Gag Q209/L210 is required for progression past the ~500S putative assembly intermediate and may serve a similar function as E207/E208 raises the possibility that a Q209-K152 hydrogen bond (Q-K hydrogen bond) in the FIV immature capsid structure could substitute for the proposed E-R salt bridge in the HIV-1 immature capsid structure and be sufficient for formation of this critical inter-hexameric helix1-helix4 CA-NTD interface. If this is the case, then the FIV Q-K hydrogen bond in place of the E-R salt bridge at the inter-hexameric CA-NTD interface could contribute to the marked instability of completed FIV immature capsids and the late (but not the early) FIV assembly intermediates relative to their HIV-1 counterparts, as shown above (Fig. 1D and 2B). While Q209 and K152 are well conserved in FIV Gag sequences from isolates obtained from domestic cats (*Felis catus*; Fig. 5D), they are not present in FIV isolates from other felines besides domestic cats with isolates from lions, cougars, bobcats, and Pallas’ cats (*Panthera leo*, *Puma concolor*, *Lynx rufus*, and *Otocolobus manul*, respectively), which more closely resemble HIV-1 at the relevant positions (Fig. 5D). This observation raises the possibility that the E-R salt bridge could form in FIV Gag isolates from lion, cougar, bobcat, or Pallas’ cat and that the immature FIV capsids formed by those isolates may be more stable than those from isolates found in domestic cats. Additionally, this might be an example of coevolved substitutions (18), since helix 1 contains an arginine when helix 4 contains a glutamic acid (e.g. in HIV-1 and FIV isolates from species besides domestic cat), while helix 1 contains a lysine when helix 4 contains a glutamine (e.g. in most FIV isolates from domestic cats). Notably, arginine forms bonds in more directions and is therefore thought to be more stabilizing than lysine (61), thus providing another reason, in addition to the salt bridge, for why the E-R pair may be more stabilizing than the Q-K pair. Future mutational studies could address whether the instability of FIV immature capsid and the ~500S assembly intermediate results from having a Q-K pair in place of the E-R pair at the inter-hexameric CA-NTD interface.

Another important HIV-1 interface is the CA-CTD dimer interface, formed by residues W316/M317 in helix 9 of HIV CA (16, 46). Our finding that mutation of the corresponding two hydrophobic residues (Y311/L312) in FIV CA helix 9 inhibited VLP production and arrested the assembly pathway at the same assembly intermediate as observed with the HIV-1 CA-CTD dimer interface mutant, argue that helix 9 in CA-CTD plays a similar role in FIV and HIV-1 immature capsid assembly, and likely forms the dimer interface in both cases. Although a previous study found that deletion of FIV CA helix 9 and part of helix 10 did not inhibit VLP production (9), deletions and point mutations can have very different phenotypes and are not always comparable.

Finally, a key implication of our findings is that compounds that target the ABCE1- and DDX6-containing RNA granules co-opted by these lentiviral Gag proteins could have broad acting antiretroviral activity, inhibiting both primate and non-primate lentiviruses and possibly other retroviruses. Notably, small molecules that target ABCE1-containing complexes are known to effectively inhibit rabies virus assembly *in vitro*, thereby providing proof-of-principle that ABCE1-containing assemblysomes are druggable (32). The finding that rabies virus, HIV-1 and other primate lentiviruses, and the non-primate lentivirus FIV, all form ABCE1-containing assemblysomes also suggests a much broader conservation of this mechanism of assembly.

## Materials and Methods

### Construction of proviral and codon-optimized FIV Gag plasmids

The proviral clone FIV 34TF10A was obtained from the NIH AIDS Reagent program (NIH AIDs Reagent Program, catalog #1236). The previously described *orfA* premature stop codon (TGA) in this proviral clone (34) was reverted to tryptophan (TGG) by overlap extension PCR. This construct was used to generate FIV, in which the FIV promoter is replaced with CMV promoter, using a previously described strategy (36). To make FIV pro-, a non-overlapping site-directed mutagenesis approach was used to introduce the D30N mutation into the FIV protease domain (numbered according to the start of protease)(62). FIV pro- env- was made by introducing two stop codons in *env* at position 7175 (numbering according to (33) – Genbank accession number: M25381.1), to generate an Env truncation mutant similar to (63). The G2A mutation was introduced into FIV pro- env- by overlap extension PCR. FIV MACA pro- env- was made by introducing two stop codons after L357 at the end of CA by overlap extension PCR. FIV CO-Gag was ordered as a gene block (Integrated DNA technologies) and cloned into pcDNA3.1/Neo(+) using HindIII and XbaI. FIV CO-Gag G2A, FIV CO-Gag P438A/P441A, FIV CO-Gag Q209A/L210A, and FIV CO-Gag Y311A/L312A were made by overlapping site-directed mutagenesis. FIV CO-Gag MACA was made by introducing a stop codon after L357 at the end of CA by overlapping site-directed mutagenesis. HIV-1 experiments utilized an HIV-1 LAI provirus that was also *pro*-, *env*-, *vpr*- and contained the *puromycin* gene in place of *nef*. To generate this construct, XhoI and XbaI were utilized to insert *puromycin* in the place of *nef* starting with HIV-GFP *env*- (27, 64), resulting in HIV env- puro. The vpr mutation was made by introducing two stop codons at the NcoI site in *vpr* by overlap extension PCR, resulting in HIV env-vpr-puro. The protease mutations (D25K, G49W, I50W) where introduced in two steps by overlap extension PCR, resulting in HIV env- vpr- pro- puro. FIV CO-Gag sequence and the sequences of the oligo-nucleotides used to generate each mutation are available upon request.

### Cells and Transfection

The feline astrocyte cell line G355-5 was obtained from ATCC (CRL-2033), and maintained in McCoy 5A (modified) medium (Life Technologies, 16600-108) supplemented with 10% fetal bovine serum (Thermo Fisher, 26140079). Cells were transfected with 1 to 6 μg DNA in 6 cm dishes using X-treameGENE 9 (Roche, 6365787001). The COS-1 cell line was obtained from ATCC (CRL-1650), and maintained in DMEM media (Life Technologies, 10566-024) supplemented with 10% fetal bovine serum (Thermo Fisher, 26140079). The 293T/17 cell line was obtained from ATCC (CRL-11268), and maintained in DMEM media (Life Technologies, 10566-024) supplemented with 10% fetal bovine serum (Thermo Fisher, 26140079). COS-1 cells were transfected with 2 μg DNA in 6 cm dishes using PEI as described previously (25). Transfection of 293T/17 cells for the proximity ligation assay is described below.

### FIV virus production and RT assay

G355-5 cells were transfected as described above but in 6-well plates. Briefly, each well was transfected with either 2 μg per well of FIV or mock transfected. At 24 hours post transfection cells were washed with media twice and media was replaced with 4 ml of complete McCoy 5A. At 144 hours post transfection media was filtered with a 0.45 micron syringe filter, aliquoted, and stored at −80C. Reverse transcriptase activity in media from FIV or mock transfected cells was estimated using SG-PERT (65). A set of recombinant RT standards (EMD Millipore catalog #382129) ranging from 5.12e3 nU/mL to 5.12e9 nU/mL was used to estimate the nU/mL of reverse transcriptase activity in the FIV stock.

### Cell lysis and DSP cross-linking

For all experiments, cells were harvested 24-48 hours post transfection. When harvested without DSP cross-linking, cells were washed twice with wash buffer (DPBS with no Calcium and no Magnesium, Thermo Fisher, 14190250) and lysates collected in the presence of protease inhibitor cocktail (Sigma Aldrich, P8340), Ribolock RNase Inhibitor (Thermo Fisher, E00381) in 1x lysis buffer (10 mM Tris pH 7.4, 100 mM NaCl, 50 mM KCl, 0.625% NP40). When harvested with DSP cross-linking verses mock treatment, cells were washed twice with wash buffer, and treated for 30 minutes either with wash buffer containing 0.1 mM DSP (Thermo Fisher, 22586) and 0.5% DMSO or treated with wash buffer with 0.5% DMSO alone (mock). DSP solution or mock treatment was removed from the cells and the reaction quenched with 1x lysis buffer without detergent for 15 minutes, and cells were subsequently harvested as described for cells without DSP crosslinking. Following harvest, cells were sheared 20 times with 18G syringe and clarified by centrifugation at slow speed (Beckman 22R, F241.5, 200xg, 10 minutes, 4C) followed by centrifugation at high speed (Beckman 22R, F241.5, 18,000xg, 1 minute, 4C).

### Virus-like particle harvest

Culture supernatants were collected 24-48 hours post transfection and filtered with 0.45 micron PES syringe filter to remove any remaining cells. Virus-like particles were centrifuged through a 30% sucrose cushion (Beckman SW55Ti, 130,000xg, 30 minutes, 4C). For WB, virus pellet was harvested directly for analysis by WB with an antibody to FIV CA-CTD (NIH AIDs Reagent Program, 4814). For velocity sedimentation, virus pellets were resuspended in the presence of protease inhibitor cocktail (Sigma Aldrich, P8340), Ribolock RNase Inhibitor (Thermo Fisher, E00381) in 1x PBS (Fisher, BP665-1). Harvested virus pellets were shaken 1000 rpm for 20 minutes at room temperature to further resuspended VLPs. Resuspended VLPs were clarified by centrifugation (Beckman 22R, F241.5, 18,000xg, 1 minute, 4C). Where indicated, resuspended virus pellets were cross-linked for 30 minutes at room temperature prior to de-enveloping by addition of DSP to a final concentration of 0. 125 mM. The cross-linking reaction was quenched with addition of Tris pH 7.4 to a final concentration of 20 mM. Virus was de-enveloped by treatment with 1/10 volume of lysis buffer with 10x NP40 detergent (10 mM Tris pH 7.4, 100 mM NaCl, 50 mM KCl, 6.25% NP40) followed by incubation on ice for 15 minutes. De-enveloped virus was clarified by centrifugation (Beckman 22R, F241.5, 18,000xg, 1 minute, 4C).

### Velocity sedimentation

Harvested virus pellet or cell lysate were layered on a step gradient (10%, 15%, 20%, 40%, 50%, 66%, and 80% sucrose in lysis buffer) and subjected to velocity sedimentation (Beckman MLS-50, 217,000xg, 45 minutes, 4°C). Gradients were fractionated from top to bottom and pelleted material was harvested for WB. Aliquots of fractions and pellet were analyzed by SDS-PAGE, followed by WB with either an antibody to FIV CA-CTD (NIH AIDs Reagent Program, 4814) or antibody to HIV CA-CTD (NIH AIDs Reagent Program, 1513), as indicated in the figure legends. Gag in pellet, which likely represents denatured Gag, was not displayed in the quantification of the velocity sedimentation gradients and represented less than 10% of total Gag signal in the gradient. The method for estimating the migration of particles with different S values in gradients has been described previously (20).

### Immunoprecipitation and western blotting

Following DSP cross-linking and cell harvest, lysates were subjected to immunoprecipitation with 1-2 μg of an antibody to ABCE1 (26), DDX6 (Bethyl, A300-461A), or rabbit IgG antibody (Bethyl, P120-101) by incubating antibody with lysate overnight at 4C with rotation, followed by incubation with protein-G coupled Dynabeads (Life Technologies, 10004D) for 4 hr with rotation. Beads were then washed twice with 1x lysis buffer and once with 1x lysis buffer without detergent. Immunoprecipitation elutes were analyzed by WB, with an antibody directed against FIV CA-CTD (NIH AIDs Reagent Program, 4814). WB signals were detected using Pierce ECL substrate (Thermo Fisher Scientific) with Carestream Kodak Biomax Light film. For detection of Gag in total cell lysates, velocity sedimentation fractions, and membrane flotation fractions, WBs were performed as described above, or using antibodies conjugated to infrared dyes (LI-COR, Lincoln, NE). Quantification of Gag bands on film was performed using Image Sutdio software (LI-COR).

### Proximity Ligation Assay (PLA)

293T/17 cells were plated into 6-well dishes containing poly-L-lysine-coated coverslips with Grace Biolabs CultureWell silicone chambers (Sigma-Aldrich) attached to create four chambers on each coverslip. Cells were transfected with 3 μg of plasmid per well using PEI as described previously (25) and 16.5 hours later were fixed for 15 minutes in 4% paraformaldehyde in PBS pH 7.4, permeabilized in 0.3% Triton X in PBS, pH 7.4 for 10 minutes, and blocked in Duolink blocking solution (Sigma-Aldrich) at 37°C for 30 min. Cells were incubated in primary antibody (described under immunoprecipitation method), followed by Duolink reagents (Sigma-Aldrich): oligo-linked secondary antibody, ligation mix, and red amplification/detection mix, with washes in between, as per the Duolink protocol. For concurrent immunofluorescence (IF), cells were incubated for 15 minutes at RT with 1:1000 Alexafluor 488 anti-mouse secondary antibody following the final 1x Buffer B PLA washes. Cover slips were mounted using Duolink In Situ Mounting Media with DAPI, sealed to the glass slides with clear nail polish, allowed to dry for 24 h at RT, and stored at −20°C. Imaging was performed with a Zeiss Axiovert 200M deconvolution microscope using Zeiss Plan-Apochromat 63X/ aperture 1.4 objective with oil immersion, with AxioVision Rel. 4.8 software. For quantification, five fields containing at least three IF-positive cells were chosen at random and imaged using identical exposure times for the red and green channels (red/green exposures were 1 second and 2.5 seconds, respectively). Images were captured as a Z-stack of five 1-μm slices centered on the focal point for the PLA. Images were deconvolved using the AxioVision software, then central Z-stack image was exported as tif files, and Image J was used to outline Gag-positive cells in each field. Within those IF positive cells, the central Z-stack image was used to count the number of PLA “spots”, and quantify IF intensity where indicated, using Image J. PLA spot number for each field was then normalized to the average IF intensity within that field, and the results were plotted with error bars representing the SEM for five fields. Prior to export of the images used in the figures, the gain on all red channels was adjusted to 3 in the AxioVision software to allow spots to be easily seen by eye. Images were then imported in 8-bit color into Adobe Illustrator to create the final figure layout, without further adjustments to color balance or gamma correction. Images shown in figures display an image from the center slice of a Z-stack that is representative of the mean. Experiment was repeated once, with similar results.

### Structure Alignment and Sequence Analysis

FIV-34TF10A Gag CA-NTD was submitted to SWISS-MODEL (66–69) which uses Blast and HHBlits to identify evolutionarily related structures matching the submitted target sequence. The top scoring template, RELIK CA-NTD (PDB accession number 2XGU.1.A; (19)) was selected for homology modeling using SWISS-MODEL. The resulting FIV CA-NTD had GMQE score of 0.73, QMEAN score of −0.17, and a normalized QMEAN4 Z-score < 1. Local quality estimates of the model were largely > 0.6. The FIV CA-NTD model obtained from SWISS-MODEL and FIV CA-CTD crystal structure (PDB accession number 5DCK;(18)) were separately aligned with HIV-1 CA structure (PDB accession number 5L93; (46)) using FATCAT (70, 71), resulting in optimized RMSD of 2.75 Å for the NTD domains and 1.94 Å for the CTD domains. The sequence alignments from FATCAT are shown in Fig. 5A and the superposed domains, rendered by CCP4MG (72), are shown in Fig. 5B.

For pairwise sequence alignments of human ABCE1 and DDX6 (Genbank accession number: NP_001035809.1 and NP_001244120.1, respectively) with feline ABE1 and DDX6 (Genbank accession number: XP_003985024.1 and XP_011284692.1, respectively), EMBOSS Needle (73) was used with default settings (73).

For FIV Gag alignments, FIV Gag sequences were obtained from NCBI protein database using the search terms “Feline immunodeficiency virus Gag” resulting in return of 618 records. These records were then filtered to select only records that at least contained Gag sequence that spanned the residues of interest (based on FIV Gag sequence in Fig. 5, this include position 152 in helix 1 to position 210 in helix 4), reducing the dataset from 618 to 405 sequences. These records were annotated for species (see Supplementary Table 1 for accession numbers and species annotation) and then aligned using Clustal Omega (73).

## Acknowledgments

The following reagents were obtained through the NIH AIDS Reagent Program, Division of AIDS, NIAID, NIH: FIV-34TF10 from Dr. John Elder; Anti-FIV p24 Monoclonal (PAK3-2C1) from DAIDS, NIAID (Custom Monoclonals, Inc.); Anti-HIV-1 p24 Hybridoma (183-H12-5C) from Dr. Bruce Chesebro and Dr. Hardy Chen. These studies were funded by Prosetta Biosciences and by NAID RO1 AI106397 to JRL. Neither of the funders (Prosetta or NIH) had a role in study design, data collection and interpretation, or the decision to submit the work for publication.

## References

1. Katrin H. 2012. Clinical Aspects of Feline Retroviruses: A Review. Viruses 4:2684–2710.

2. Elizabeth WU, Marcus M, James KC, Janet KY. 2008. Advances in FIV vaccine technology. Veterinary Immunology and Immunopathology 123:65–80.

3. Dunham SP, Bruce J, MacKay S, Golder M, Jarrett O, Neil JC. 2006. Limited efficacy of an inactivated feline immunodeficiency virus vaccine. Veterinary Record 158:561–562.

4. Levy J, Crawford C, Hartmann K, Hofmann-Lehmann R, Little S, Sundahl E, Thayer V. 2008. 2008 American Association of Feline Practitioners’ feline retrovirus management guidelines. J Feline Medicine Surg 10.

5. Katrin H, Anita W, Michèle B. 2015. Efficacy of Antiviral Drugs against Feline Immunodeficiency Virus. Veterinary Sciences 2:456–476.

6. Dorothee B. 2014. FIV in cats-a useful model of HIV in people? Veterinary Immunology and Immunopathology 159:171–179.

7. Freed EO. 2015. HIV-1 assembly, release and maturation. Nat Rev Microbiol 13:484–496.

8. Mariana LM, Cristina C, Silvia AG, José LA. 2001. Mutational analysis of the feline immunodeficiency virus matrix protein. Virus Research 76:103–113.

9. Abdusetir Cerfoglio JC, González SA, Affranchino JLL. 2014. Structural elements in the Gag polyprotein of feline immunodeficiency virus involved in Gag self-association and assembly. The Journal of general virology 95:2050–2059.

10. Mariana LM, MarÍa LR, Silvia AG, José LA. 2004. Functional domains in the feline immunodeficiency virus nucleocapsid protein. Virology 327:83–92.

11. Benjamin GL, Miranda S-X, Dimiter GD, Catherine SA, Ferri S, Kunio N, Andrew GS, Robert JF, Eric OF. 2008. Molecular Characterization of Feline Immunodeficiency Virus Budding. Journal of Virology 82:2106–2119.

12. Elder JH, Schnölzer M, Hasselkus-Light CS, Henson M, Lerner DA, Phillips TR, Wagaman PC, Kent SB. 1993. Identification of proteolytic processing sites within the Gag and Pol polyproteins of feline immunodeficiency virus. Journal of virology 67:1869–1876.

13. Lingappa JR, Reed JC, Tanaka M, Chutiraka K, Robinson BA. 2014. How HIV-1 Gag assembles in cells: Putting together pieces of the puzzle. Virus Res doi:10.1016/j.virusres.2014.07.001.

14. Patarca R, Haseltine WA. 1985. A major retroviral core protein related to EPA and TIMP. Nature 318:390.

15. Orlinsky KJ, Gu J, Hoyt M, Sandmeyer S, Menees TM. 1996. Mutations in the Ty3 major homology region affect multiple steps in Ty3 retrotransposition. J Virol 70:3440–3448.

16. Gamble TR, Yoo S, Vajdos FF, von Schwedler UK, Worthylake DK, Wang H, McCutcheon JP, Sundquist WI, Hill CP. 1997. Structure of the carboxyl-terminal dimerization domain of the HIV-1 capsid protein. Science 278:849–853.

17. Schur FK, Hagen WJ, Rumlova M, Ruml T, Muller B, Krausslich HG, Briggs JA. 2015. Structure of the immature HIV-1 capsid in intact virus particles at 8.8 A resolution. Nature 517:505–508.

18. Khwaja A, Galilee M, Marx A, Alian A. 2016. Structure of FIV capsid C-terminal domain demonstrates lentiviral evasion of genetic fragility by coevolved substitutions. Sci Rep 6:24957.

19. Goldstone DC, Yap MW, Robertson LE, Haire LF, Taylor WR, Katzourakis A, Stoye JP, Taylor IA. 2010. Structural and functional analysis of prehistoric lentiviruses uncovers an ancient molecular interface. Cell Host Microbe 8:248–259.

20. Lingappa JR, Hill RL, Wong ML, Hegde RS. 1997. A multistep, ATP-dependent pathway for assembly of human immunodeficiency virus capsids in a cell-free system. J Cell Biol 136:567–581.

21. Dooher JE, Schneider BL, Reed JC, Lingappa JR. 2007. Host ABCE1 is at Plasma Membrane HIV Assembly Sites and Its Dissociation from Gag is Linked to Subsequent Events of Virus Production. Traffic 8:195–211.

22. Robinson BA, Reed JC, Geary CD, Swain JV, Lingappa JR. 2014. A temporospatial map that defines specific steps at which critical surfaces in the Gag MA and CA domains act during immature HIV-1 capsid assembly in cells. J Virol 88:5718–5741.

23. Lingappa JR, Dooher JE, Newman MA, Kiser PK, Klein KC. 2006. Basic residues in the nucleocapsid domain of Gag are required for interaction of HIV-1 gag with ABCE1 (HP68), a cellular protein important for HIV-1 capsid assembly. J Biol Chem 281:3773–3784.

24. Klein KC, Reed JC, Tanaka M, Nguyen VT, Giri S, Lingappa JR. 2011. HIV Gag-leucine zipper chimeras form ABCE1-containing intermediates and RNase-resistant immature capsids similar to those formed by wild-type HIV-1 Gag. J Virol 85:7419–7435.

25. Tanaka M, Robinson BA, Chutiraka K, Geary CD, Reed JC, Lingappa JR. 2015. Mutations of Conserved Residues in the Major Homology Region Arrest Assembling HIV-1 Gag as a Membrane-Targeted Intermediate Containing Genomic RNA and Cellular Proteins. J Virol 90:1944–1963.

26. Zimmerman C, Klein KC, Kiser PK, Singh ARS, Firestein BL, Riba SC, Lingappa JR. 2002. Identification of a host protein essential for assembly of immature HIV-1 capsids. Nature 415:88–92.

27. Reed JC, Molter B, Geary CD, McNevin J, McElrath J, Giri S, Klein KC, Lingappa JR. 2012. HIV-1 Gag co-opts a cellular complex containing DDX6, a helicase that facilitates capsid assembly. J Cell Biol 198:439–456.

28. Engeland CE, Brown NP, Borner K, Schumann M, Krause E, Kaderali L, Muller GA, Krausslich HG. 2014. Proteome analysis of the HIV-1 Gag interactome. Virology 460–461:194–206.

29. Kedersha N, Anderson P. 2007. Mammalian stress granules and processing bodies. Methods Enzymol 431:61–81.

30. Ostareck DH, Naarmann-de Vries IS, Ostareck-Lederer A. 2014. DDX6 and its orthologs as modulators of cellular and viral RNA expression. Wiley Interdiscip Rev RNA 5:659–678.

31. Dooher JE, Lingappa JR. 2004. Conservation of a step-wise, energy-sensitive pathway involving HP68 for assembly of primate lentiviral capsids in cells. J Virol 78:1645–1656.

32. Lingappa UF, Wu X, Macieik A, Yu SF, Atuegbu A, Corpuz M, Francis J, Nichols C, Calayag A, Shi H, Ellison JA, Harrell EK, Asundi V, Lingappa JR, Prasad MD, Lipkin WI, Dey D, Hurt CR, Lingappa VR, Hansen WJ, Rupprecht CE. 2013. Host-rabies virus protein-protein interactions as draggable antiviral targets. Proceedings of the National Academy of Sciences of the United States of America 110:E861–868.

33. Talbott RL, Sparger EE, Lovelace KM, Fitch WM, Pedersen NC, Luciw PA, Elder JH. 1989. Nucleotide sequence and genomic organization of feline immunodeficiency virus. Proceedings of the National Academy of Sciences of the United States of America 86:5743–5747.

34. Waters AK, De Parseval AP, Lerner DL, Neil JC, Thompson FJ, Elder JH. 1996. Influence of ORF2 on host cell tropism of feline immunodeficiency virus. Virology 215:10–16.

35. Hind JF, Dyana TS, Eric MP. 2012. Construction and Testing of orfA +/− FIV Reporter Viruses. Viruses 4:184–199.

36. Poeschla EM, Wong-Staal F, Looney DJ. 1998. Efficient transduction of nondividing humancells by feline immunodeficiency virus lentiviral vectors. Efficient transduction of nondividing human cells by feline immunodeficiency virus lentiviral vectors 4.

37. Lin Y-C, Brik A, Parseval Ad, Tam K, Torbett BE, Wong C-H, Elder JH. 2006. Altered Gag Polyprotein Cleavage Specificity of Feline Immunodeficiency Virus/Human Immunodeficiency Virus Mutant Proteases as Demonstrated in a Cell-Based Expression System. Altered Gag Polyprotein Cleavage Specificity of Feline Immunodeficiency Virus/Human Immunodeficiency Virus Mutant Proteases as Demonstrated in a Cell-Based Expression System 80.

38. Gottlinger HG, Dorfman T, Sodroski JG, Haseltine WA. 1991. Effect of mutations affectingthe p6 gag protein on human immunodeficiency virus particle release. Proc Natl Acad Sci U S A 88:3195–3199.

39. Huang M, Orenstein JM, Martin MA, Freed EO. 1995. p6Gag is required for particleproduction from full-length human immunodeficiency virus type 1 molecular clones expressingprotease. J Virol 69:6810–6818.

40. Demirov DG, Orenstein JM, Freed EO. 2002. The late domain of human immunodeficiencyvirus type 1 p6 promotes virus release in a cell type-dependent manner. J Virol 76:105–117.

41. Chu HH, Chang YF, Wang CT. 2006. Mutations in the alpha-helix directly C-terminal to themajor homology region of human immunodeficiency virus type 1 capsid protein disrupt Gagmultimerization and markedly impair virus particle production. J Biomed Sci 13:645–656.

42. Joshi A, Nagashima K, Freed EO. 2006. Mutation of dileucine-like motifs in the humanimmunodeficiency virus type 1 capsid disrupts gag-virus assembly, gag interactions, gag-membrane binding, and virion maturation. J Virol 80:7939–7951.

43. Ono A, Waheed AA, Joshi A, Freed EO. 2005. Association of human immunodeficiency virustype 1 gag with membrane does not require highly basic sequences in the nucleocapsid: use of anovel Gag multimerization assay. J Virol 79:14131–14140.

44. von Schwedler UK, Stray KM, Garrus JE, Sundquist WI. 2003. Functional surfaces of thehuman immunodeficiency virus type 1 capsid protein. J Virol 77:5439–5450.

45. Grover JR, Llewellyn GN, Soheilian F, Nagashima K, Veatch SL, Ono A. 2013. Roles playedby capsid-dependent induction of membrane curvature and Gag-ESCRT interactions in tetherinrecruitment to HIV-1 assembly sites. J Virol 87:4650–4664.

46. Schur FK, Obr M, Hagen WJ, Wan W, Jakobi AJ, Kirkpatrick JM, Sachse C, Krausslich HG, Briggs JA. 2016. An atomic model of HIV-1 capsid-SP1 reveals structures regulatingassembly and maturation. Science 353:506–508.

47. Esteva MJJ, Affranchino JLL, González SA. 2014. Lentiviral Gag assembly analyzed through the functional characterization of chimeric simian immunodeficiency viruses expressing different domains of the feline immunodeficiency virus capsid protein. PloS one 9.

48. Soderberg O, Gullberg M, Jarvius M, Ridderstrale K, Leuchowius KJ, Jarvius J, Wester K, Hydbring P, Bahram F, Larsson LG, Landegren U. 2006. Direct observation of individual endogenous protein complexes in situ by proximity ligation. Nat Methods 3:995–1000.

49. Anderson P, Kedersha N. 2006. RNA granules. J Cell Biol 172:803–808.

50. Beckham CJ, Parker R. 2008. P bodies, stress granules, and viral life cycles. Cell Host Microbe 3:206–212.

51. Eulalio A, Behm-Ansmant I, Izaurralde E. 2007. P bodies: at the crossroads of post-transcriptional pathways. Nat Rev Mol Cell Biol 8:9–22.

52. Verschoor A, Warner JR, Srivastava S, Grassucci RA, Frank J. 1998. Three-dimensional structure of the yeast ribosome. Nucleic Acids Res 26:655–661.

53. Chatel-Chaix L, Clement JF, Martel C, Beriault V, Gatignol A, DesGroseillers L, Mouland AJ. 2004. Identification of Staufen in the human immunodeficiency virus type 1 Gag ribonucleoprotein complex and a role in generating infectious viral particles. Mol Cell Biol 24:2637–2648.

54. Abrahamyan LG, Chatel-Chaix L, Ajamian L, Milev MP, Monette A, Clement JF, Song R, Lehmann M, DesGroseillers L, Laughrea M, Boccaccio G, Mouland AJ. 2010. Novel Staufen1 ribonucleoproteins prevent formation of stress granules but favour encapsidation of HIV-1 genomic RNA. J Cell Sci 123:369–383.

55. Bouttier M, Saumet A, Peter M, Courgnaud V, Schmidt U, Cazevieille C, Bertrand E, Lecellier CH. 2011. Retroviral GAG proteins recruit AGO2 on viral RNAs without affecting RNA accumulation and translation. Nucleic acids research doi:10.1093/nar/gkr762.

56. Izumi T, Burdick R, Shigemi M, Plisov S, Hu WS, Pathak VK. 2013. Mov10 and APOBEC3G localization to processing bodies is not required for virion incorporation and antiviral activity. J Virol 87:11047–11062.

57. Cinti A, Le Sage V, Ghanem M, Mouland AJ. 2016. HIV-1 Gag Blocks Selenite-Induced Stress Granule Assembly by Altering the mRNA Cap-Binding Complex. MBio 7:e00329.

58. Valiente-Echeverria F, Melnychuk L, Vyboh K, Ajamian L, Gallouzi IE, Bernard N, Mouland AJ. 2014. eEF2 and Ras-GAP SH3 domain-binding protein (G3BP1) modulate stress granule assembly during HIV-1 infection. Nat Commun 5:4819.

59. Phalora PK, Sherer NM, Wolinsky SM, Swanson CM, Malim MH. 2012. HIV-1 replication and APOBEC3 antiviral activity are not regulated by P bodies. J Virol 86:11712–11724.

60. Decker CJ, Parker R. 2012. P-bodies and stress granules: possible roles in the control of translation and mRNA degradation. Cold Spring Harb Perspect Biol 4:a012286.

61. Sokalingam S, Raghunathan G, Soundrarajan N, Lee SG. 2012. A study on the effect of surface lysine to arginine mutagenesis on protein stability and structure using green fluorescent protein. PLoS One 7:e40410.

62. Laco GS, Schalk-Hihi C, Lubkowski J, Morris G, Zdanov A, Olson A, Elder JH, Wlodawer A, Gustchina A. 1997. Crystal structures of the inactive D30N mutant of feline immunodeficiency virus protease complexed with a substrate and an inhibitor. Biochemistry 36:10696–10708.

63. Kemler I, Barraza R, Poeschla EM. 2002. Mapping the encapsidation determinants of feline immunodeficiency virus. J Virol 76:11889–11903.

64. Yamashita M, Emerman M. 2004. Capsid is a dominant determinant of retrovirus infectivity in nondividing cells. J Virol 78:5670–5678.

65. Vermeire J, Naessens E, Vanderstraeten H, Landi A, Iannucci V, Van Nuffel A, Taghon T, Pizzato M, Verhasselt B. 2012. Quantification of reverse transcriptase activity by real-time PCR as a fast and accurate method for titration of HIV, lenti-and retroviral vectors. PLoS One 7:e50859.

66. Biasini M, Bienert S, Waterhouse A, Arnold K, Studer G, Schmidt T, Kiefer F, Gallo Cassarino T, Bertoni M, Bordoli L, Schwede T. 2014. SWISS-MODEL: modelling protein tertiary and quaternary structure using evolutionary information. Nucleic Acids Res 42:W252–258.

67. Kiefer F, Arnold K, Kunzli M, Bordoli L, Schwede T. 2009. The SWISS-MODEL Repository and associated resources. Nucleic Acids Res 37:D387–392.

68. Arnold K, Bordoli L, Kopp J, Schwede T. 2006. The SWISS-MODEL workspace: a web-based environment for protein structure homology modelling. Bioinformatics 22:195–201.

69. Guex N, Peitsch MC, Schwede T. 2009. Automated comparative protein structure modeling with SWISS-MODEL and Swiss-PdbViewer: a historical perspective. Electrophoresis 30 Suppl 1:S162–173.

70. Ye Y, Godzik A. 2003. Flexible structure alignment by chaining aligned fragment pairs allowing twists. Bioinformatics 19 Suppl 2:ii246–255.

71. Ye Y, Godzik A. 2004. FATCAT: a web server for flexible structure comparison and structure similarity searching. Nucleic Acids Res 32:W582–585.

72. Winn MD, Ballard CC, Cowtan KD, Dodson EJ, Emsley P, Evans PR, Keegan RM, Krissinel EB, Leslie AG, McCoy A, McNicholas SJ, Murshudov GN, Pannu NS, Potterton EA, Powell HR, Read RJ, Vagin A, Wilson KS. 2011. Overview of the CCP4 suite and current developments. Acta Crystallogr D Biol Crystallogr 67:235–242.

73. Li W, Cowley A, Uludag M, Gur T, McWilliam H, Squizzato S, Park YM, Buso N, Lopez R. 2015. The EMBL-EBI bioinformatics web and programmatic tools framework. Nucleic Acids Res 43:W580–584.

